# Estimation of chloroplast macromolecular complex copy numbers and subunit stoichiometries during the *Chlamydomonas reinhardtii* cell cycle

**DOI:** 10.64898/2026.03.30.715394

**Authors:** Stefan Schmollinger, Daniela Strenkert, Samuel O. Purvine, Carrie D. Nicora, Eric Soubeyrand, Gilles J. Basset, Sabeeha S. Merchant

## Abstract

An unbiased, quantitative view of biomolecules in a living cell is a prerequisite for accurate modeling approaches and informs our understanding of cellular metabolism at scale. In this work, we used the total protein approach (TPA), in which the total protein mass of a given proteomics sample is used as a calibrator for absolute protein quantification, to determine protein abundances during the *Chlamydomonas reinhardtii* diurnal cycle. We use external, independently measured quantitative markers (metals, pigments) to assess the absolute protein abundances in unlabeled whole cell extracts. We calculate protein abundances in fg / cell of 7322 Chlamydomonas proteins, 2266 of which were captured in every time point, including the major proteins involved in the light reactions, photoprotection, proteostasis and fatty acid metabolism during a cell cycle. As expected, Rubisco large and small subunits are present in a 1:1 stoichiometry, with the large subunit being the most abundant protein in our data set, averaging 5.05 × 10^6^ molecules per cell, reflecting 2.7% of the total protein mass. We noticed that PSII is the most abundant complex involved in the light reactions with 2.08 × 10^6^ complexes per cell. PSI averages 1.75 × 10^6^ complexes per cell and cytochrome *b*_6_*f* averages 0.77 × 10^6^ complexes per cell. The TPA is a robust tool to study proteome dynamics quantitatively, while avoiding artefacts due to biochemical fractionation. Our proteome data set with an unprecedented temporal resolution is a valuable resource to assess protein abundances during the cell cycle in the reference alga Chlamydomonas.

## INTRODUCTION

*Chlamydomonas reinhardtii* (Chlamydomonas hereafter) is a chlorophyte, single-celled alga, sharing fundamental metabolic pathways with other eukaryotic algae and most importantly land plants. Hence, it has been a valuable reference organism, especially for studies of eukaryotic photosynthesis and chloroplast biology (1, 2). Besides a chromosome-level, high-quality assembly of the nuclear and plastid genomes with structural and functional annotations, decades of experimental work with Chlamydomonas has generated an impressive set of molecular tools, which have been exploited for discoveries on acclimation responses, the cell cycle, photosynthesis and cilium structure and function (3–9).

An important but challenging aspect of genome-wide biology is the quantitative assessment of biomolecule concentrations in a cell, which is also a prerequisite for accurate mathematical modeling approaches (10). Prior to the development of mass spectrometry methods, protein copy numbers and stoichiometries of subunits in complexes were determined individually by spectroscopic methods for photosystem components or immunochemistry for well-studied proteins (11–13). Nevertheless, these methods are labor-intensive to scale to the entire proteome, and quantitative proteomics has been utilized instead to allow for the untargeted identification and quantitation of a large set of proteins in a sample (14). Accurate MS-based protein quantification is challenging and can roughly be divided into label-free and label-based techniques. Label-free methods rely on features directly associated with the measurement, especially the number of identification events in a run (unique peptides/spectral counts) or quantitative features of the identification event (peak area/height from an associated liquid chromatography run), while label-based experiments involve an additional labelling step, often the isotopic modification of proteins or peptides, either as part of the experimental workflow or with the addition of external standards. Both approaches can be useful for either relative or, by the use of standards, absolute protein quantification at a larger scale. Label-based methods include metabolic labelling of the proteome with ^15^N or ^13^C (via labelled amino acids or alternatively the N source) during cell growth (15), the addition of isobaric TAGs to peptides during MS sample preparation (16) or the use of quantification concatamers (QconCAT) (17), all of which have been applied to distinguish the abundance of proteins in Chlamydomonas (18–21). These approaches offer high sensitivity and quantitative precision even in relatively complex samples, but they can be expensive and they further increase the complexity of the analyte. Labelling approaches can result in reduced discovery or they add additional measurement time. Therefore, there remains considerable interest in label-free methodologies, which offer greater protein identification in complex samples and do not require additional processing during sample preparation that comes at the cost of lower accuracy.

Label-free absolute quantitation of proteins has been used successfully for describing the proteomes of several organisms, including yeast, human and bacteria (22). While there are many approaches to determine absolute protein abundances in untargeted proteomics, the more accurate workflows rely on quantitative features (peak area/height) of the measurements and employ the known concentration of a class of native proteins present in cell extracts, or the calculated total protein mass, as internal calibrators to yield protein copy numbers / cell. In the case of the proteomic ruler approach (23, 24), the MS signal of histones is used as an internal standard in a proteomics experiment, since the amount of histones is proportional to the amount of DNA present in a cell. The exact amount of DNA per cell is known for many species (genome size x ploidy) or can be determined experimentally. The total protein approach (TPA) is another label free method for absolute quantitation of proteins. The method uses the summed total MS signal from all identified proteins in a proteomics sample to reflect the total protein mass in a cell, using the rational assumption that the most abundant proteins in a cell determine its bulk protein content (25). The total protein mass of a given sample can be assessed on a per cell basis by protein concentration assays and cell counting. This method can be used to quantify thousands of proteins per sample (25–27).

We used the TPA for Chlamydomonas in previous work to assess protein abundances in cells grown with various amounts of Zn supplementation (28). In that work, we considered the TPA superior to the proteomic ruler approach, as we noted that abundances of histones were regularly underestimated in these proteomic data sets. We assumed that this is due to their extensive posttranslational modifications and the large number and non-ideal distribution of lysines and arginines in the histone primary sequence, restricting the number of tryptic peptides detected. We therefore use the label free TPA in this work to assess copy numbers of proteins involved in photosynthesis and carbon fixation and to determine changes in protein abundances and stoichiometry adjustments as cells progress through the cell cycle.

The Chlamydomonas cell cycle progresses over a 24h period that is set by a diurnal light regime, typically 12h light and 12h dark (29–31). Starting with time 0 at the onset of light, there is a series of macromolecular metabolic events as cells grow in the G1 phase, followed by DNA synthesis (S phase), mitosis(M) early in the dark phase, cytokinesis followed by a G0 phase. While RNA-sequencing technologies enabled complete quantitation of changes in the transcriptomes, including plastid and other non-polyadenylated RNAs, during the Chlamydomonas cell cycle (30, 31), complementary data are still lacking for the proteome.

Here, we show that the total protein ruler method is suitable for absolute quantifications of a large set of Chlamydomonas proteins. The power of this study is frequent sampling throughout the cell cycle (15 time points) in experimental triplicates (derived from independent photobioreactors) under conditions where cells double exactly once on average. We use high resolution mass spectrometry and peak intensity-based protein quantification (MASIC) combined with the TPA to measure relative and absolute abundances (in fg/cell) of ∼7322 Chlamydomonas proteins. We combine these data with copy number estimates of dozens of proteins and calculate subunit stoichiometries in multi-subunit complexes, with a focus on participants in the light reactions.

## RESULTS

### Proteome profiling of the Chlamydomonas diurnal cycle

Recent technological advances in mass spectrometry (MS)-based proteomics have further facilitated large-scale protein detection and quantitation. In this study, we aimed for an organism-wide analysis of protein dynamics during the diurnal cycle using a label-free approach on unfractionated, total cell lysates. By doing so, our fractionation-free method reduces the risk of introducing artificial changes in protein abundances that result from changes in solubility whilst enabling higher sample throughput. To this end, we collected 16 samples (in triplicate) from highly synchronized cells that were grown over a full 24-hour diurnal cycle in independent flat panel photobioreactors as described previously (30), with one additional time point collected during the beginning of the following night (+13) (Figure 1 and Supplemental Figure 1). Experiments were highly reproducible between the three independent photobioreactor runs, and optical densities (as proxy for biomass) increased as cells grew and accumulated biomolecules during the light phase, ultimately resulting in exactly two daughter cells at the beginning of the following night (Supplemental Figure 1). In total we identified 50,074 unique peptides in our proteomic dataset, associated with 10743 proteins, 7322 of which were identified with at least two unique peptides (Figure 1a, Supplemental Figure 2, Supplemental Table 1).

**Figure 1.**
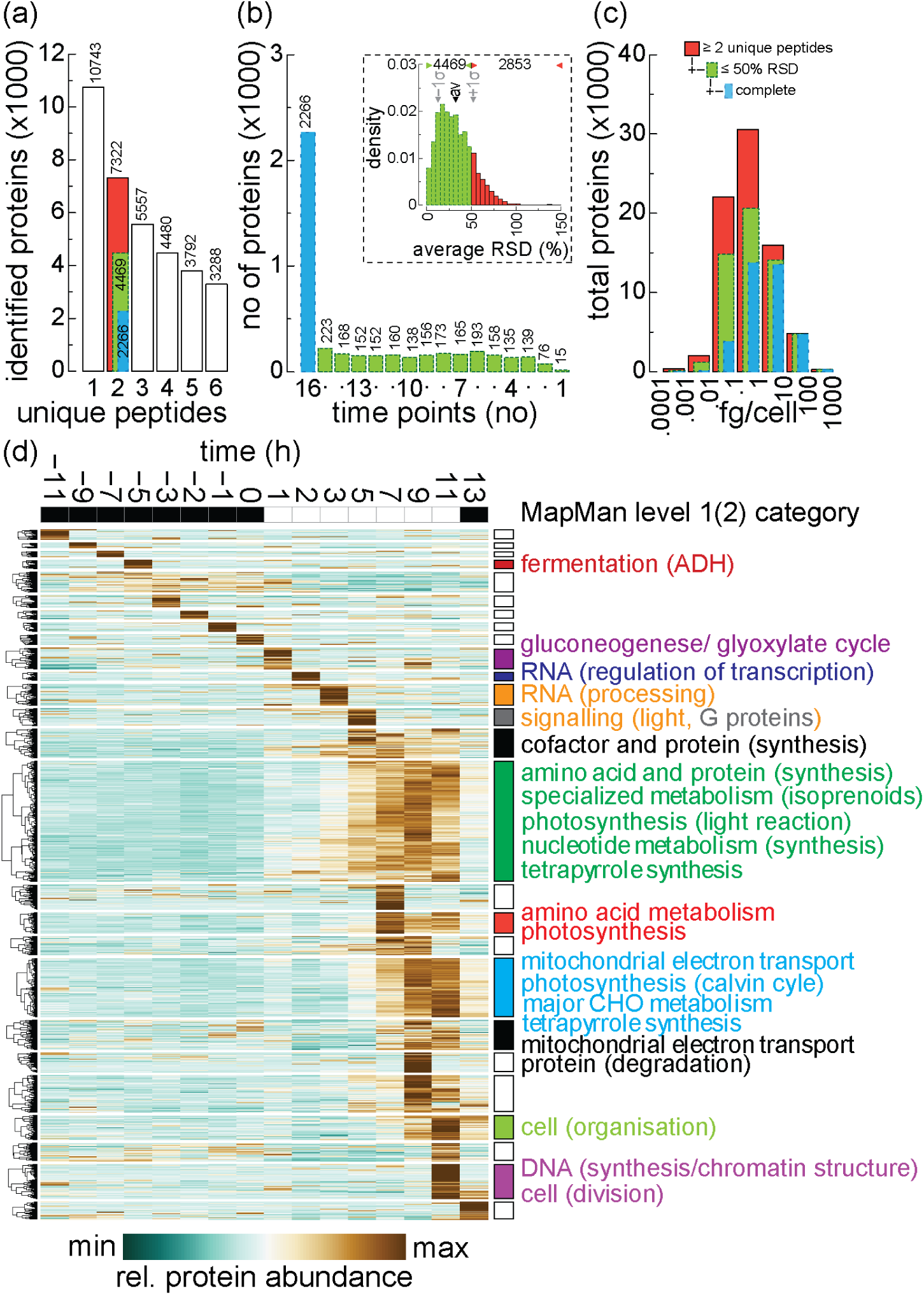
The Chlamydomonas proteome is highly rhythmic over the diurnal cycle. (a) Number of proteins identified in the dataset dependent on the number of required unique peptides for each protein. The used cutoff (≥2 unique peptides, red fill) is highlighted, including the fraction with an average relative standard deviation (RSD) below 50% (green fill, blue in complete series). (b) Length of the timeseries for the 4469 identified proteins with an average RSD < 50%. The complete series’ are again highlighted in blue. The distribution of the average RSD is shown in the inset (top right), average, +/− one standard deviation and cutoff are highlighted by arrows. (c) Histogram of the distribution of absolute protein abundances (fg/cell) for proteins identified with ≥2 unique peptides (red fill) and the subset identified with an av. RSD < 50% (green fill, blue fill for complete series). (d) Heatmap representation of rel. protein abundances for 4469 proteins (≥2 unique peptides, av. RSD <50%) over one diurnal cycle. Proteins were clustered hierarchically and organized by peak abundance into 26 groups, level 1 MapMan categories enriched in a respective cluster (α < 0.01) are indicated on the right side of the cluster, enriched level 2 MapMan categories are included in brackets (α < 0.01). Protein abundances were standardized (z-score) and n.d. values were replaced with the lowest abundance determined for any protein in the dataset.

### Absolute protein abundances per cell using the total protein ruler method

MASIC was used for peak-based peptide quantification, yielding relative protein abundances (32). In order to estimate absolute protein abundances and determine protein copy numbers per cell, the total summed mass spectrometry signal of all measured peptides (reflecting the cumulative protein mass) correlates with the measured protein abundance in each sample, allowing absolute protein abundance estimates. This method has the advantage that it does not rely on the accurate quantification of individual specific proteins for calibration nor requires the utilization of isotopically distinguished standards, either at the protein or peptide level. Accordingly, we determined the cellular protein content for each sample along the diurnal cycle, which we estimated to be between 14 pg / cell during the night and 38 pg / cell during the day (Supplemental Figure 1). The protein per cell estimates were in the same range as the estimated average of ∼21 pg protein / cell in non-synchronous, photoheterotrophically-grown Chlamydomonas cultures, estimated by us and others (18, 28, 34). As is customary, we used multiple different peptides of each protein in a sample to determine average protein abundances and subsequently calculated relative standard deviations (RSD) of protein abundances between independent replicates. We found an average RSD across timepoints of 32% for all proteins in the dataset (Figure 1b, inlet). To remove less reliably quantified proteins from further analysis we required an average RSD across timepoints to be below 50%, which was equivalent to the average RSD +1 standard deviation, and simultaneously required confident protein identification in at least two of the three replicates of each time point of the diurnal cycle. We found a total of 4469 proteins (out of 7322, ∼ 61%) matching those criteria (high confidence dataset), 2266 of which were identified in each point of the time course (∼ 51% of the high confidence dataset, complete series, Figure 1a, b), while only 15 proteins (∼ 0.3%) could be reliably identified in just a single time point (Figure 1b). We observed protein abundance changes spanning ∼6 order of magnitudes, with more abundant proteins generally showing a lower variance and a greater likelihood of being recovered throughout the time course (Figure 1c, Supplemental Figure 2). The individual TPA calibrated protein abundances showed a median increase of 2.9-fold along the diurnal cycle (Supplemental Figure 2b), equivalent to the measured cellular protein content, which was used for scaling, indicating that the used transformations accurately reflected the dataset as a whole. The vast majority of identified and reliably quantified proteins doubled in abundance during the light phase of the cell cycle (clusters 10-25, 3685 proteins, ∼82 %), consistent with the cell’s productive phase and the need to double its protein inventory to accommodate a doubling event in this experimental setup (Figure 1d, Supplemental Figure 1, 2b). We used a hierarchical clustering approach on the high confidence data set (4469 proteins) to organize the data, yielding a total of 26 distinct clusters (Figure 1, Supplemental Table 2). Automatic and manual functional annotations were used in the MapMan gene ontology framework to identify biological processes enriched in these clusters (http://pathways.mcdb.ucla.edu/algal/index.html), revealing a multitude of different pathways to be enriched at different times during the diurnal cycle (Figure 1d). Proteins involved in various fermentation processes were significantly overrepresented in cluster 4 that peaks during the night phase, while proteins involved in photosynthesis and carbon fixation were enriched in various clusters peaking during the middle of the day. Proteins involved in transcriptional regulation, light signaling, RNA processing and cofactor and protein biosynthesis were enriched in clusters with peak abundances early during the day, preceding the clusters containing the highly abundant structural proteins required for light and carbon capture. Clusters containing proteins with peak expression towards the end of the day, as cells prepared for cell division, were overrepresented for proteins involved in cell organization, DNA replication and cell division (cohesion and condensin, histone genes).

### Cuproprotein abundance as a feature enabling unbiased assessment of protein quantification accuracy using the TPA

To assess the general accuracy of the total protein ruler-based method for total protein abundance determination we sought completely independent, quantitative measurements to query protein abundance measurements. We compared the total protein-bound copper (Cu) to the total amount of intracellular Cu that was measured independently by ICP-MS/MS throughout the diurnal cycle. In Cu replete conditions, the Cu quota of the cell is largely dependent on the abundance of three major Cu proteins, namely plastocyanin, Cyt oxidase, and a multicopper ferroxidase, FOX1, which is involved in high-affinity Fe assimilation and is more abundant when Fe is limiting in the growth medium (35, 36). Phototrophically-grown asynchronous Chlamydomonas cultures were shown to contain approximately 5 × 10^6^ Cu atoms / cell (37), which was consistent with our observations ranging from 2 × 10^6^ Cu atoms / cell during the night and up to 5.5 × 10^6^ Cu atoms / cell by the end of the day. In an Fe-replete environment, such as in this diurnal cycle, ferroxidase abundance is low, and we expected the protein-bound Cu quota to be largely determined by just plastocyanin and Cyt oxidase, which was indeed the case (Figure 2). Most importantly, we noted strikingly comparable accumulation kinetics between our proteomics-based estimates of protein-bound Cu (Figure 2, stacked bars) and the estimates of total intracellular Cu by ICP-MS/MS (Figure 2, cyan line, open circles). The copper bound to plastocyanin and Cyt oxidase accounted for ∼80-90% of the total cellular Cu content, depending on the particular time point of the diurnal cycle. The protein-associated Cu slightly but consistently underestimated the total cellular Cu content, whose increase preceded the increase in abundances of the two cuproproteins (Figure 2). The lower abundance of protein-bound Cu can be attributed to the pipeline not capturing i) lower-abundance cupro-proteins that add to the remainder and ii) contributions of non-protein-bound Cu to the total Cu quota in Chlamydomonas, either contained in storage sites (acidocalcisomes) (38, 39) or bound to chelators like glutathionine, y-glutamylcysteine and cysteine (28). Both of these Cu pools were included in the ICP-MS/MS-based elemental analyses but cannot be assessed via proteomics. Nonetheless, the close match between the intracellular Cu and the abundance of cuproproteins, both in abundance and accumulation kinetics, support the use of the total protein ruler method for determining protein copy numbers from proteomics data, if certain parameters are considered – as further outlined below.

**Figure 2.**
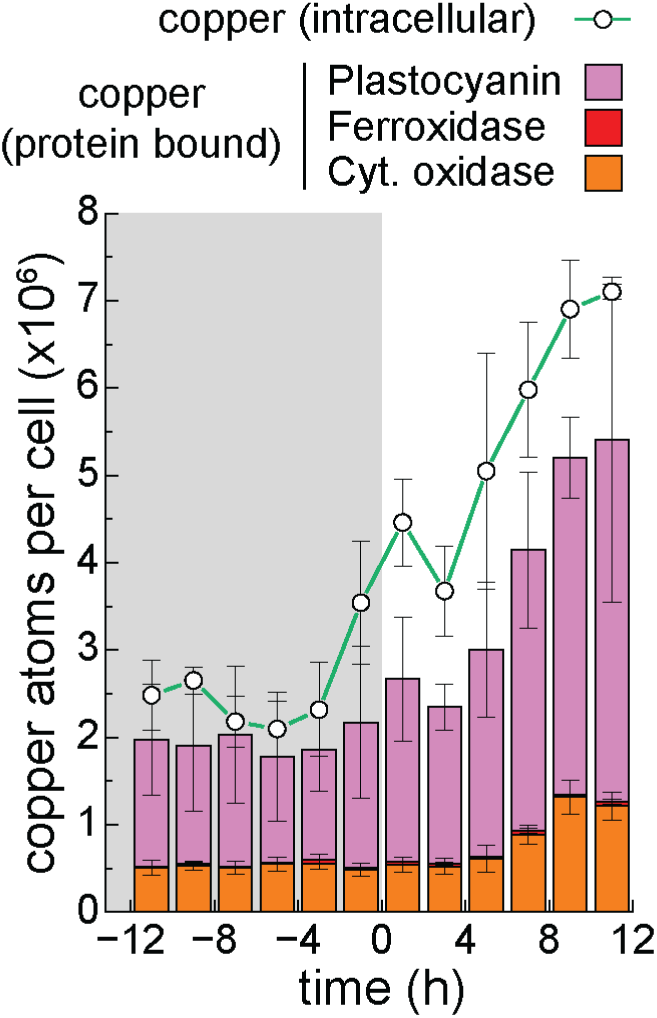
Cellular copper content correlates well with the combined abundances of the major cupro-protein throughout the diurnal cycle. The total intracellular copper content (green line, in units of millions of atoms per cell) of Chlamydomonas cultures at the indicated time points during the diurnal cycle was determined by ICP-MS/MS, shown are averages and standard deviation between three independent bioreactors. The protein-bound copper content in the major cellular copper proteins (stacked bars; plastocyanin in pink, ferroxidase in red, cytochrome oxidase in orange) was estimated by quantitative proteomics using the proteomic ruler method, taking into account the number of copper binding sites in each protein to yield the total amount of bound copper in millions of atoms per cell. Cells in triplicate bioreactors were grown in iron-replete conditions, where ferroxidase abundance is low.

### RuBisCO as proof of principle for multi-subunit stoichiometry determination

We next analyzed the composition of the total protein pool recovered during the diurnal cycle. The most abundant proteins made up a large fraction of the total cellular protein content, the top 20 most abundant proteins combined constituted more than 20% and the top 100 proteins made up almost half (∼44%) of the total measured protein mass despite representing only a small fraction of the number of proteins detected in the dataset (100 out of 10743, 0.9%, Figure 3a). We noted that the top 1000 proteins accounted for almost the entire proteome mass (86%), suggesting that, while coverage of individual proteins in our proteomics data set is not yet exhaustive, we capture almost all proteins that represent the protein mass in a Chlamydomonas cell. As expected for phototrophically-grown cultures supplemented with low/air levels of CO_2_ (∼0.04%), structural subunits of complexes involved in photosynthesis and the CO_2_ assimilation machinery were the biggest constituency among the most abundant proteins in the dataset. They were reliably recovered throughout the diurnal cycle and their induction during the light phase was consistent with their functional role. The most abundant protein in the dataset was indeed the large subunit of ribulose-1,5-bisphosphate carboxylase / oxygenase (RuBisCO), which catalyzes the carboxylation of the substrate ribulose-1,5-bisphosphate, the first step in CO_2_ fixation by the Calvin-Benson cycle. Form I RuBisCO, found in green algae and vascular plants, is a hexadecamer composed of eight large plastid encoded subunits (RbcL) and eight small nucleus-encoded subunits (RbcS) (Figure 3b, c). RbcL and RbcS average 450 and 126 fg / cell in our data, respectively, reflecting 2.7% and 0.7% of the total protein mass, which individually place both proteins in the top 20 most abundant proteins based on mass %. Copy number estimates for both RuBisCO subunits revealed average abundances of 5.05 ×10^6^ for RbcS and 4.93 ×10^6^ for RbcL subunits per cell, which is in line with the expected stoichiometry between the large and the small subunit (our observed average ratio is 1:1.004). The stoichiometry between both subunits also was remarkable consistent throughout the diurnal cycle. In addition, we estimate that the cells contain an average of 6.24 ×10^5^ hexadecameric RuBisCOs per cell during the diurnal period, with peak abundances reaching an impressive 10.8 ×10^5^ RuBisCO molecules per cell by the end of the light phase. This abundance is comparable, albeit lower, than earlier proteomics reports ((Table 1) and (18)), which could be a consequence of differences in growth conditions (phototrophic diurnal growth vs. photoheterotrophic growth in continuous light), strain choice or methodology employed in each of these studies. Taken together, abundant soluble protein complexes, as exemplified by RuBisCO, were recovered stoichiometrically along the diurnal cycle.

**Figure 3.**
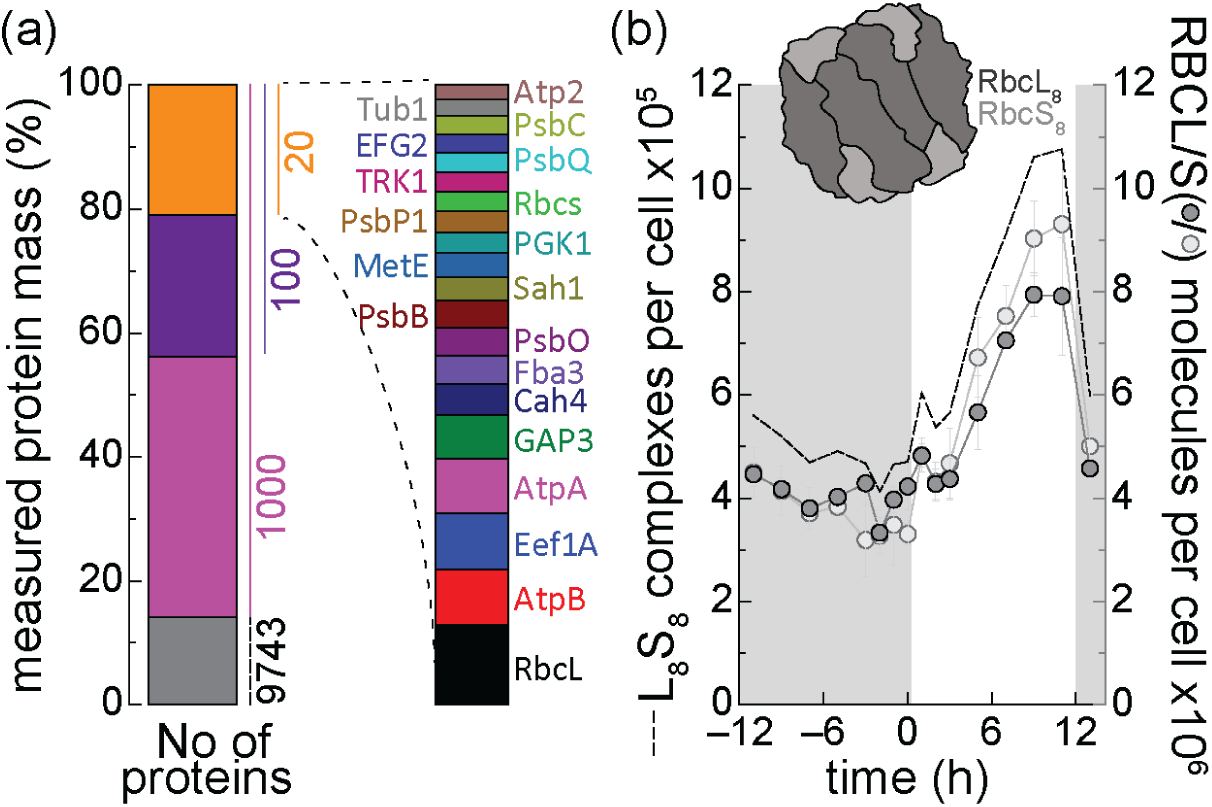
Protein composition of Chlamydomonas: (a) Contribution (%) of the top 20 / 100 / 1000 most abundant proteins of Chlamydomonas to the total protein mass among all observed proteins; the contribution of each of the top 20 proteins is further highlighted on the right. (b) Abundances of RbcL and RbcS in units of millions of molecules per cell, and the total RuBisCO complexes (L_8_S_8_) per cell were estimated by quantitative proteomics over the diurnal cycle using the proteomic ruler method.

**Table 1.**

|  | Number of protein subunits per cell<br>x 10<sup>6</sup> |  |  |  |  |  | Ratio between complexes |  |
| --- | --- | --- | --- | --- | --- | --- | --- | --- |
|  | PS2 | PS1 | Cyt<br>*b<sub>6</sub>f* | RbcL | RBCS | PCY | PS2:PS1:Cyt<br>*b<sub>6</sub>f* | RbcL:RBCS |
| This study<sup>a</sup> | 2.1 | 1.8 | 0.8 | 5.1 | 5 | 2.2 | 2.7:2.27:1 | 1.004:1 |
| Hammel et al.<br>2018 | 3.1 | 1.8 | 1.7 | 15 | 10 | 2.1 | 1.8:1.1:1 | 1.56:1 |
| Wienkoop et<br>al. 2010 | 1.4-2.5 |  | 0.9-1.6 | 20-26 | 0.6-1.7 | 1-5 | 9-29:n:1 | 11-44:1 |
| Murakami et<br>al. 1997 |  |  |  |  |  |  | 1.9:0.6:1 |  |
| Chow et al.<br>1990 |  |  |  |  |  |  | 2.7:1.4:1 |  |

<sup>a</sup>the respective averages of all time points during the cell cycle were used to compare to other studies.

### A quantitative perspective on the light reactions

#### Complexes and electron carriers of the chloroplast electron transfer chain

Four abundant membrane protein complexes in the thylakoid membrane are photosystems II (PSII) and I (PSI) and the cytochrome (Cyt) *b*_6_*f* complex, which catalyze the transfer of electrons from water through the plastoquinone (PQ) pool to ferredoxin (Fd), concomitant with vectorial trans-thylakoid membrane H^+^ transfer and the ATP synthase, which uses the H^+^ gradient to drive ATP synthesis. Reduced Fd is a source of electrons for many plastid metabolic pathways either directly or indirectly though NADPH production by Fd NADP+ oxidoreductase (Figure 4, top, visualized according to (40)).

**Figure 4.**
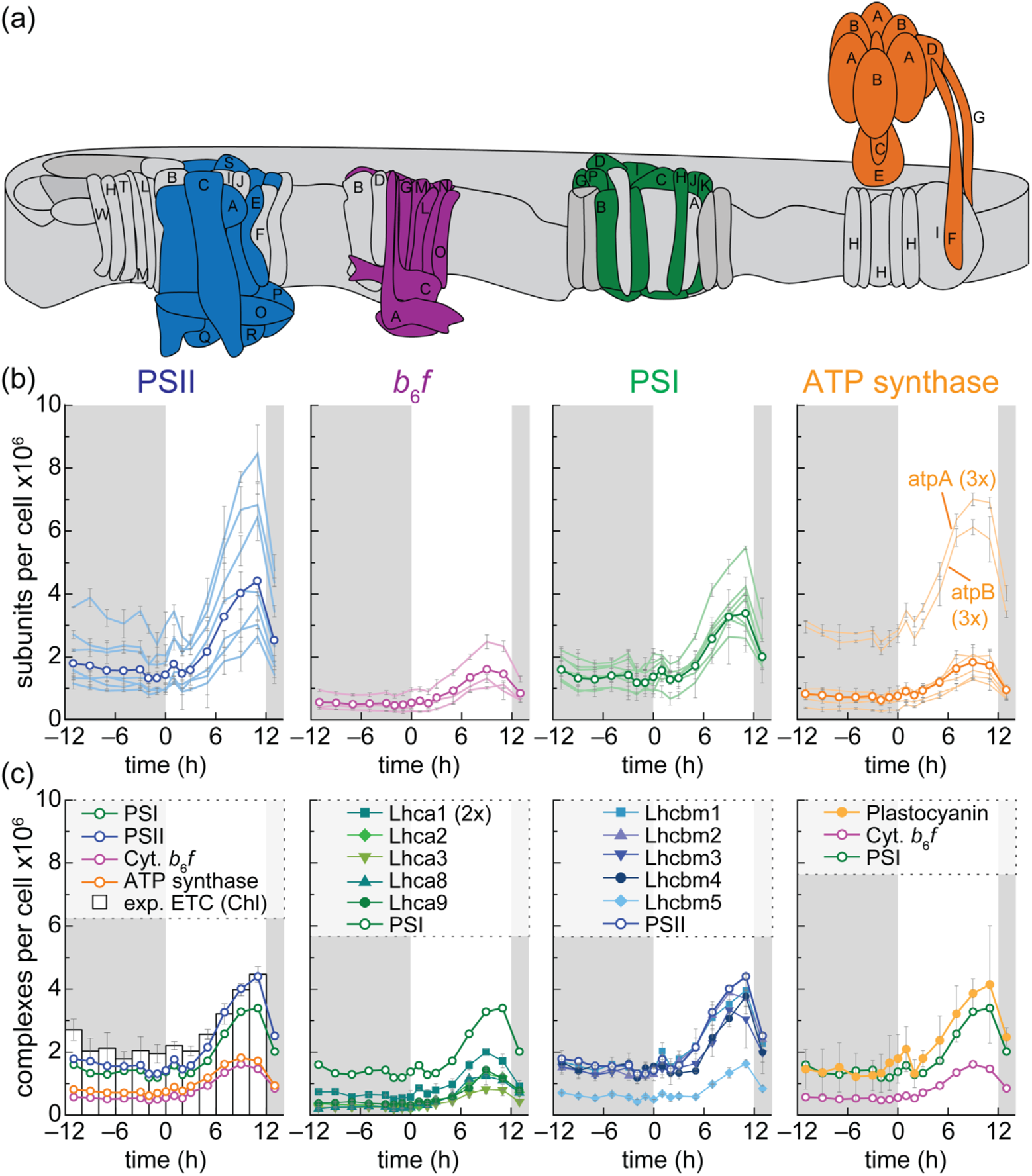
Absolute protein abundances and stoichiometry changes of subunits within the chloroplast electron transfer chain during the cell cycle. (a) Schematics depicting the four major complexes of the photosynthetic ETC, subunits that weren’t quantitatively recovered are shown in grey. (b) protein abundance of each subunit (in units of millions per cell) was estimated by quantitative proteomics using the proteomic ruler method. (c) Amount of each of the complexes per cell, compared to the expected amounts calculated from Chl extracts over the cell cycle. Individual complex abundances related to individual antenna proteins (Lhca and Lhcbm) and electron carriers (Plastocyanin).

The relative abundances of PSII and PSI are important for maintaining flux under changing light conditions, and for balancing linear vs. cyclic electron flow to accommodate changes in NADPH vs. ATP demand. The unique photochemical properties of these complexes have allowed absolute quantification of photosystems by spectroscopy and accordingly there is considerable literature on photosystem abundance and ratios of PSII to PSI in various organisms under various environmental conditions (41–44). Quantification of the Cyt *b*_6_*f* complex by spectroscopy is more challenging because of the smaller number of chromophores per complex and hence lower signals, and for the ATP synthase it is not possible in the absence of spectroscopic signatures (45). The advent of genomics enabled the use of mass spectrometry-based methods for estimating protein abundances (18), but the outcomes indicated a range of values in the literature for individual proteins of the same complex (Table 1). Quantification of proteins by mass spectrometry depends not only on their abundances but also on the reliable generation of peptides and their retrieval and capture, which depends on the physical properties of the protein; especially solubility, hydrophobicity, number and accessibility of protease cleavage sites – for membrane proteins, typically low and non-randomly distributed along their length. Hydrophobicity is a key consideration because a) photosystem proteins (e.g. D1, D2, PsaA/B) can be especially hydrophobic, and b) it limits the observability of a peptide and hence quantification of the corresponding protein. There are fewer Lys and Arg residues in these proteins, hence fewer tryptic sites, and fewer peptides. The Lys and Arg residues when they do occur may be close together in regions exposed to the solvent, like extra-membrane loops, perhaps disproportionately distributed on one side of the membrane, resulting in very small peptides that also evade observation. There are typically also fewer side chains that are amenable to ionization in the trans-membrane regions, leading to fewer spectral signals relative to more polar peptides. Therefore, we used the GRAVY (Grand Average of Hydropathy) index, a measure of hydrophobicity (46), for assessing the reliability of quantification of individual Psb, Psa, Atp, Pet and Lhc proteins for our estimation of the stoichiometries of components of the photosynthetic apparatus in the thylakoid membrane (Figure 4, top). The average GRAVY index was calculated for the entirety of the experimentally quantified peptide sequence that was captured by mass spectrometry for a given protein and only polypeptides with an average negative GRAVY index (indicative for hydrophilic regions) were considered for abundance assessment of the respective complex (Supplemental Table 3).

For each complex, we averaged the estimated abundance of each such reliably quantified polypeptide, taking into account subunit stoichiometry of α and β for the ATP synthase, to arrive at the copy number per cell of each of the four major complexes of the photosynthetic apparatus (Figure 4a, Supplemental Table 3). As expected, subunits of all four major complexes (PSII, Cyt *b*_6_*f*, PSI, ATP synthase) were tightly co-expressed during the cell cycle, with PSI and PSII averaging 1.75 × 10^6^ and 2.08 × 10^6^ complexes per cell (Figure 4 and Table 1). Cyt *b*_6_*f* and ATPase complexes were found to be sub-stoichiometric (∼45%) relative to the photosystem core complexes, averaging ∼ 7.7 × 10^5^ and 9.8 × 10^5^ complexes per cell, a phenomenon that has been observed previously for Cyt *b*_6_*f* relative to PSI by others (Table 1). Eukaryotic algae use chlorophyll (Chl) *a* and *b* for light harvesting and charge separation in their photosynthetic reaction centers. Chl *a* and *b* are protein-bound, can be extracted and accurately quantified using spectroscopic methods (47), allowing us to assess absolute protein abundances of the highly abundant photosynthetic complexes in phototrophic organisms. We used absolute quantitative Chl *a* and *b* abundance measurements from our previous study to calculate the expected amount of photosynthetic electron transfer chains at each time point during the diurnal cycle (Figure 4c, bars), assuming state 1, a 1:1 stoichiometry between PSI and PSII, a Chl *a* to *b* ratio of 2.6:1, CP26/29 and 3.5 LHCII trimers/PSII (48) and found that the amounts closely resemble the observed, average protein abundances and accumulation dynamics of PSI and PSII core complexes found in the mass spectrometry dataset. Based on Chl measurements we expect an average of ∼ 2.6 × 10^6^ ETCs per cell.

Each Chlamydomonas photosystem is associated with accessory antenna of Chl-binding proteins. The LHCI complex associated with PSI consists of the Lhca subunits, Lhca1 through Lhca9. Lhca2 and Lhca9 are associated as a heterodimer on one side of PSI, while the others form two layers of tetrameric, half-moon shaped belts on the other side: Lhca1, Lhca8, Lhca7 and Lhca3 occur in an inner belt from PsaG to PsaK, and Lhca1, Lhca4, Lhca6 and Lhca5 in an outer belt (49). Upon transition to state 2, two trimers of phosphorylated Lhcbm proteins also associate with PSI through PsaO to shift excitation energy input into the photosystems towards PSI (50).

Lhca1 was present at 9.05 × 10^5^ molecules per cell on average, which represents a ∼ 1.9-fold higher abundance as compared to Lhca2, Lhca3, Lhca8 and Lhca9, which averaged 5.09 × 10^5^, 3.73 × 10^5^, 4.85 × 10^5^ and 5.65 × 10^5^ molecules / cell, respectively. Lhca3 and Lhca8 were ∼20% lower in abundance on average as compared to Lhca2 and Lhca9. Consistent with previous reports based on biochemical and structural work (50), we capture the 2:1 stoichiometry of Lhca1 relative to the other inner belt Lhca subunits. We note also that Lhca proteins appear to be sub-stoichiometric to PSI perhaps because we do not fully recover them quantitatively((Figure 4) and (49, 51)).

The protein products of the *LHCB* and *LHCBM* genes associate with the PSII reaction center (RC) to form the PSII-LHCII super-complexes. The protein inventory includes Lhcb4, Lhcb5 and Lhcb7, but transcriptomic and proteomic data suggest that only the *LHCB4* and *LHCB5* genes are expressed to any appreciable extent, and hence are the only ones relevant in this proteomic study (52). In Chlamydomonas, the Lhcb4 subunit or CP29 and Lhcb5 subunit or CP26 are found in one monomeric copy adjacent to each reaction center monomer. There are 9 *LHCBM* genes (1 through 9) encoding corresponding proteins that form the trimeric LHCII complexes that associate with PSII through interactions with the monomeric Lhcb subunits. The proteins have been classified into four groups based on sequence relationships: type I (Lhcbm3, 4, 6, 8 and 9), type II (Lhcbm5), type III (Lhcb2 and 7, which are identical in sequence and hence not distinguished in proteomics experiments) and type IV (Lhcbm1) (53, 54). Expression profiles indicate different patterns and levels of expression, suggesting distinct physiological functions although the individual proteins are biochemically similar with respect to pigment, chemical and physical properties. Lhcbm1, Lhcbm2/7 and Lhcbm3 are most abundant in thylakoid membranes, and indeed structural analyses of isolated PSII-LHCII super-complexes with large antenna contained substantial amounts of each of these proteins, especially Lhcbm3 subunits (55–57). The quantity and quality of light was held constant in our experiments and these Lhcbm abundances can serve as a baseline for assessing the potential for compositional changes in LHCIIs in response to variation in light input or other environmental variables. For instance, Lhcbm9 with a low S content is typically very weakly expressed, but strongly induced in S deficiency and may have a specific role in photoprotection under stress (58). Most of the Lhcbm4/6/8 subunits are found free in the membrane, poorly connected to PSII, but they are still required for physiological LHCII abundance and state transitions (59). We noted that Lhcbm1, Lchbm2, Lhcbm3 and surprisingly also Lhcbm4 were found in stoichiometric amounts to each other and to PSII, averaging 2.02 × 10^6^, 1.93 × 10^6^, 1.73 × 10^6^ and 1.81 × 10^6^ molecules per cell (Figure 4). As expected, and based on previous studies, Lhcbm5, Lhcbm8 and Lhcbm9 are present in lower abundances relative to Lhcbm1,2,3: averaging 7.38 × 10^5^, 3.70 × 10^4^ and 3.49 × 10^4^ molecules per cell. While overall subunit abundances between Lhcbm1,2,3 and 4 were similar, the increase of Lhcbm3 and Lchbm4 subunits during the day was delayed by ∼2 h as compared to the other subunits. The fact that we capture Lhcbm4 in stoichiometric quantities with Lhcbm1,2,3 and other PSII subunits in this work whereas not in prior works could be explained by potentially weaker association of Lhcbm4 with PSII that might result in its under-representation in fractionated samples.

#### Proteins and molecules involved in photoprotection

In previous work we showed that genes encoding proteins involved in photoprotection, specifically LHCSR3, LHCSR1 and PSBS (60, 61), are up-regulated immediately after lights on, early in the day during the Chlamydomonas diurnal cycle (52). Two genes encode identical proteins, both for PSBS and LHCSR3. LHCSR3 and LHCSR1 associate with PSII and are involved in energy-dependent quenching (qE) of excess light, as part of the cell’s repertoire of non-photochemical quenching (NPQ) mechanisms (62). Both proteins were tightly co-expressed, but sub-stoichiometric to the photosystem core subunits - averaging ∼3 × 10^5^ molecules per cell (Figure 5a), ∼ 1 LHCSR protein / 6-7 PSII/LHCII complexes. Similarly, PSBS is involved in qE in land plants and assumed to be associated with PSII. While most proteins doubled during the diurnal cycle (Supplemental Figure 2b), PSBS did not. PSBS accumulated only transiently during the diel period, averaging only 2.4 × 10^4^ molecules per cell at peak abundance at 1 h after lights on, before it was degraded below detection limit by +5h in the light. Note that PSBS degradation was evident already by +2h (Figure 5a), which is consistent with our previous study using immunodetections (30). The abundance of PSBS was ∼100 times lower than that of PSII, even at peak accumulation, consistent with a catalytic or regulatory role. We note that PSBS quantification was largely based on one unique peptide with a positive GRAVY index of 0.18, suggesting that we may be underestimating its abundance in this study.

**Figure 5.**
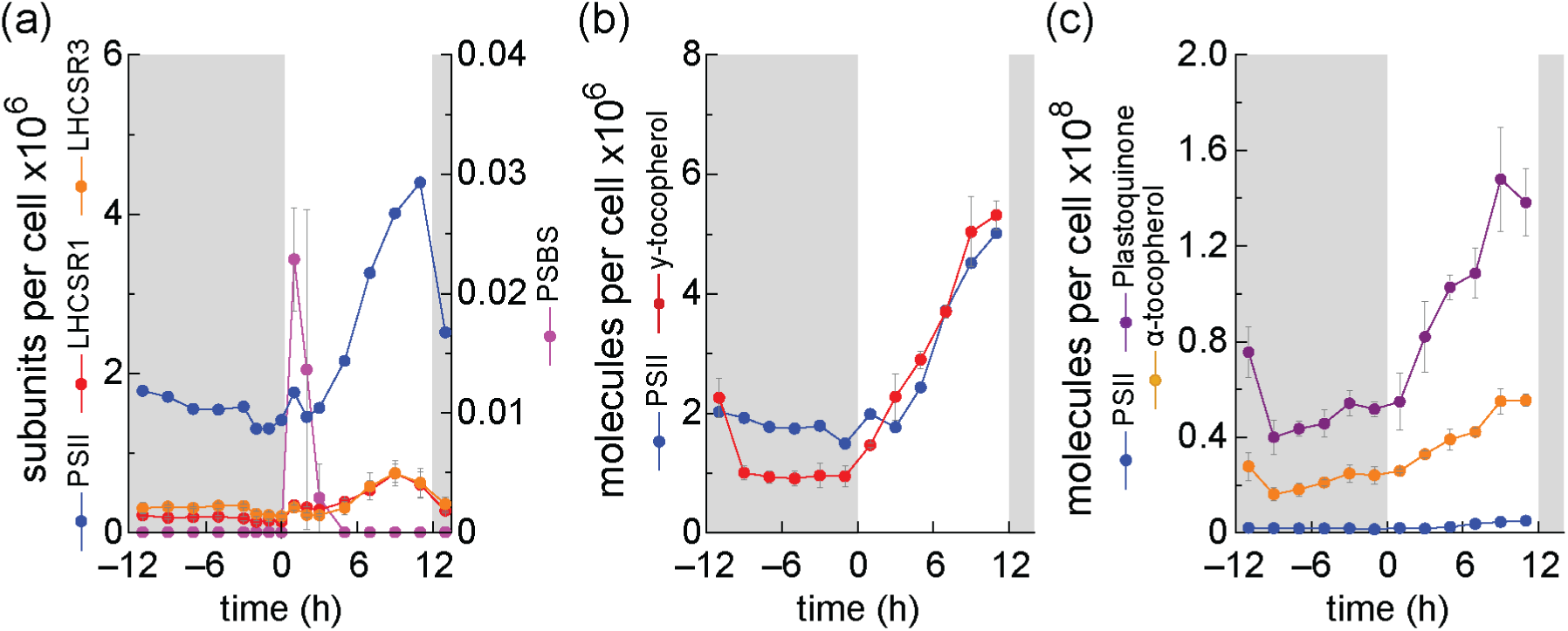
Copy number estimates of proteins and molecules involved in photoprotection. The abundance of each protein or molecule (in units of millions of molecules per cell) was estimated by quantitative proteomics using the proteomic ruler method (LHCSR1, LHCSR3, PSBS) or by HPLC (α- and y-tocopherols, PQ).

Besides the protein components of the photosystems, thylakoid membranes contain redox active lipids, namely plastoquinone-9 (PQ), which is a reactant in the electron transport chain, and tocopherols, which have anti-oxidant function especially in a situation of over-excitation of PSII and over-reduction of the PQ pool. In previous work (30), we quantified the content and redox state of PQ for chemical analyses. For this work, we used the same approach to quantify α- and γ-tocopherols and to estimate absolute abundances per cell of these critical lipid molecules. We note that γ-tocopherol is stoichiometrically matched to PSII at 2.3 × 10^6^ molecules / cell (Figure 5b) while both PQ and α-tocopherol are present in vastly excess amounts, averaging 30 and 15 times more molecules per cell as there are PSII core complexes (Figure 5c). The presence of 93% α-tocopherol vs. 7% γ -tocopherol in our work is consistent with the tocopherol pool composition observed by others in land plants and in Chlamydomonas that consist of ∼ 90% α-tocopherol and only relatively small amounts of γ-tocopherol (63–65).

#### Subunit composition of chloroplast proteases responsible for proteostasis

FtsH and ClpP proteases are proteolytic enzymes whose activities have been implicated in plastid protein quality control, photosystem subunit biosynthesis and PSII repair. ClpP is a protease consisting of two heptameric rings. The subunit composition in each ring and the masses of the mature protein of each subunit have been previously determined (66, 67). The top ring consists of one ClpR6 subunit, three ClpP4 and three ClpP5 subunits, while the bottom ring contains three ClpP1C subunits and one of the ClpR1,2,3 and 4 subunits each. Indeed, for the bottom ring, we estimate a subunit composition of 11.6 × 10^4^ ClpP1C, 4.01 × 10^4^ ClpR1, 3.7 × 10^4^ ClpR2 subunits, 5.5 × 10^4^ ClpR3 and 2.78 × 10^4^ ClpR4 subunits per cell reflecting a 3-1.04-0.96-1.4-0.7 subunit composition, which nicely matches the expected 3-1-1-1-1 arrangement. On the other hand, for the top ring, we estimate 3.02 × 10^4^ ClpR6, 1.02 × 10^5^ ClpP4 and 1.52 × 10^5^ ClpP5 subunits per cell for 1-3.8-5 stoichiometry for ClpR6-ClpP4-ClpP5 instead of the expected 1-3-3 distribution. We estimate the abundance of the individual rings to be 3.84 × 10^4^ bottom rings and 3.97 × 10^4^ top rings per cell, consistent with a 1:1 stoichiometry of each ring, which argues against uneven recovery of subunits of the top ring. The unexpected 1-3.8-5 pattern for the top ring therefore suggests *in vivo* heterogeneity with respect to the ratios of the P4 and P5 polypeptides in the composition of the top ring evident at the whole cell level (Figure 6). The absolute abundance of Clp proteases works out to be ∼ 4 × 10^4^ molecules / cell.

**Figure 6.**
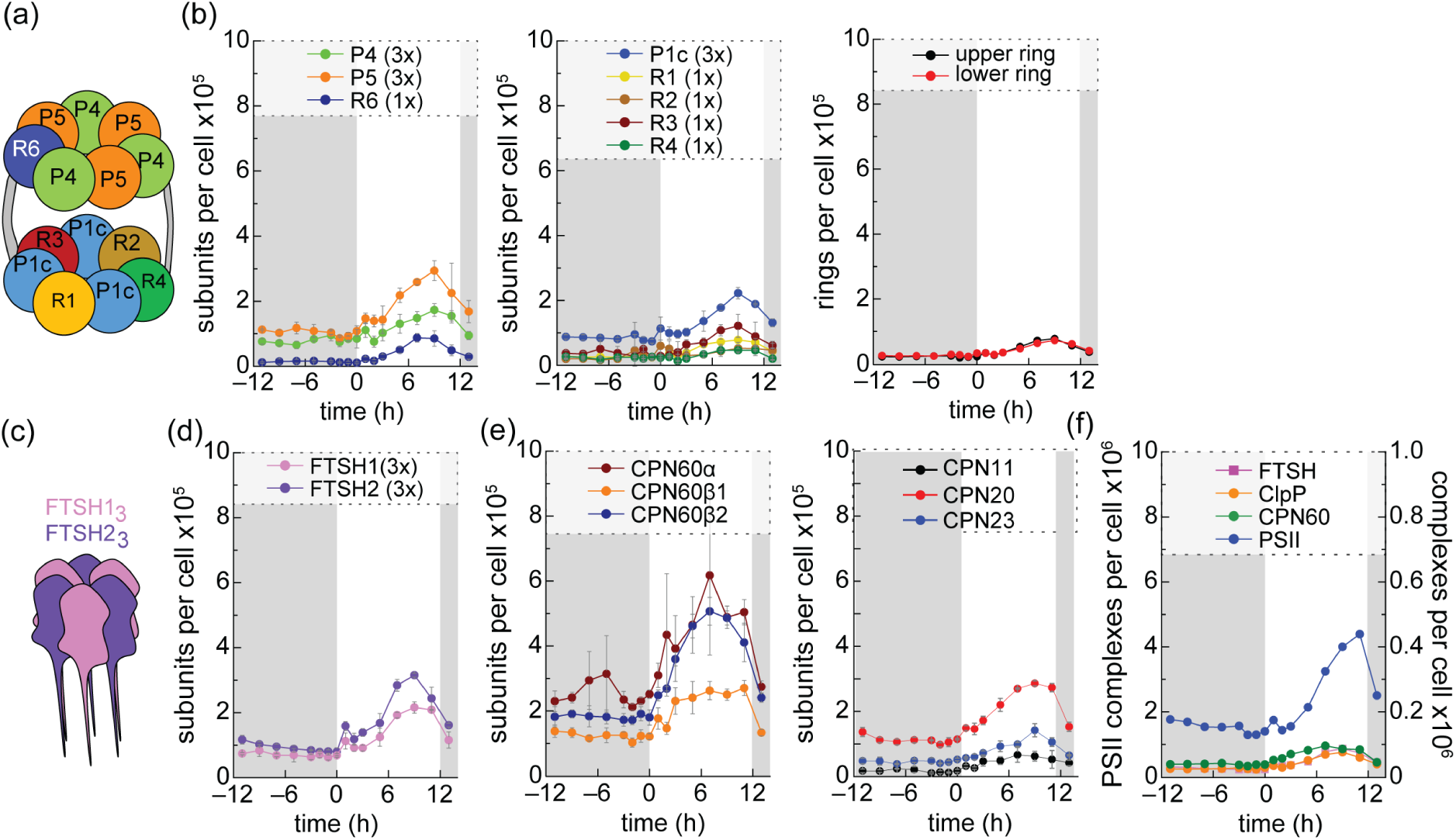
Copy number estimates of chloroplast ATP-dependent multimeric proteases. (a,c) Schematic showing a cartoon depicting the ClpP and FTSH complex and subunit composition, respectively. (b,d) the abundance of each protein (in units of millions of molecules per cell) was estimated by quantitative proteomics using the proteomic ruler method and used to (f) calculate the amount of each of the complexes (ClpP, CPN60 and FTSH) per cell over the cell cycle and how they relate to photosystem complex abundances (PSII).

FtsH is a ubiquitous protease found in bacteria and bacteria-derived organelles like mitochondria and chloroplasts. In chloroplasts, FtsH is responsible for the degradation of membrane proteins, especially the photosynthetic complexes in the thylakoid membrane (68–70). The bacterial FtsH protein is a hexameric ring of individual FtsH subunits. The organellar enzymes are similar in structure except that the chloroplast FtsH enzymes are encoded by multigene families resulting in heteromeric complexes consisting of type A and type B subunits, both of which are required, in 1:1 stoichiometric abundance, for assembly of the holoenzyme (71, 72). Chlamydomonas contains one type A and one type B subunit (Figure 6). When we estimated the abundance of each subunit type across the 24-hour diel period, we noted coordinate expression with a constant 1:1.35 ratio, close to the expected 1:1 ratio between the type A and type B subunits (Figure 6). FtsH1 averages ∼ 1.1 × 10^5^ molecules per cell and FTSH2 ∼ 1.4 × 10^5^ molecules per cell for an approximate holoenzyme abundance of ∼ 4 × 10^4^ molecules per cell. This number, which is similar to the abundance of the ClpP complexes in the chloroplast, is compatible with a catalytic role of the proteases in degrading thylakoid membrane complexes that are each found at a stoichiometry of ∼ 10^6^ per cell or about 2 orders of magnitude greater (Figure 6).

It has been shown previously that the chloroplast chaperonin system is crucial for Rubisco folding and assembly. In Chlamydomonas. The chloroplast chaperonin is formed by two hetero-olimeric, heptameric, stacked rings consisting of CPN60α, CPN60β1 and CPN60β2 subunits. The stoichiometry of chaperonin subunits (CPN60αβ1β2) was previously determined using QconCAT proteins as internal standards on fractionated cells. This approach yielded heterogenous preparations and it was estimated that the ratios between individual subunits lies between 5:3:6 and 6:2:6 for CPN60α:CPN60β1:CPN60β2 (73). We estimate that the subunit stoichiometry is 6:3:5 CPN60α:CPN60β1:CPN60β2 (based on an estimated average ratio of 6:2.96:4.85) (Figure 6). Co-chaperonins CPN11, CPN20 and CPN23 show a stoichiometry of 0.55:2.79:1.2 in our dataset, implying subunit heterogeneity. While the chaperonin complex is present at an average of 5.65 × 10^4^ complexes per cell, which is in the same range as plastid proteases, expression of chaperonin is up-regulated earlier in the day (at time 1h) as compared to Ftsh or ClpP expression (time 5h) (Figure 6). This result is consistent with chaperonin’s role in Rubisco folding, which is needed during the early hours of the day for efficient carbon assimilation, while the plastid proteases are needed towards the end of the day for protein quality control.

#### The curious case of the ACCase complex

In Chlamydomonas (as in land plants), the heteromeric acetyl CoA carboxylase (ACCase) in the chloroplast catalyzes the first step in fatty acid biosynthesis by generating malonyl CoA. The enzyme has three functionalities found in four subunits: the biotin carboxylase (BC), which activates the substrate CO_2_ in an ATP-dependent reaction, attaching it to biotin bound on biotin carboxyl carrier protein (BCCP), from where it is transferred via the carboxylase transferase (CT, made up of α and β subunits) to acetyl CoA (Figure 7) (74–76). The enzyme is a key regulatory point for fatty acid biosynthesis with activity being stimulated alkaline pH, Mg^2+^, conditions that prevail in the light when reductant for anabolic pathways is available (77). Of the four ACCase subunits, three are stoichiometric (average of 1.07 × 10^5^ molecules per cell), but BC is found at ∼3-4-fold excess (∼ 3.9 × 10^5^ per cell) (Figure 7). Despite the well-established operation of ACCase in the light (vs. the dark), per cell subunit stoichiometries are unchanged throughout the diel period, underscoring the relevance of post-translational regulation. One important regulator is biotin attachment domain containing (BADC). The BADC protein is an inactive homolog of BCCP, which can also bind to BC but lacks the conserved lysine residue required for function. In Arabidopsis, there are three BADC proteins that are typically more abundant than BC itself throughout plant development (78, 79). We identified a single algal BADC-like protein that harbors all the characteristics of an inactive BCCP protein, encoded at Cre01.g037850 and find that it too is stoichiometric with the other subunits (CT α and β subunits). Conditions that favor BADC binding to BC (e.g. at low pH in the dark) would sequester BC, blocking access ofBCCP to BC and thus reducing flux through ACCase (77). The mechanism elegantly enables a dynamic response of ACCase to the metabolic state of the chloroplast. The greater abundance of BC vs. the three other ACCase subunits is thus rationalized on the basis of competition between BCCP and BADC for binding to BC, When fatty acid synthesis is required, the over-abundance of BC will ensure that it can associate with BCCP and form an active ACCase complex.

**Figure 7.**
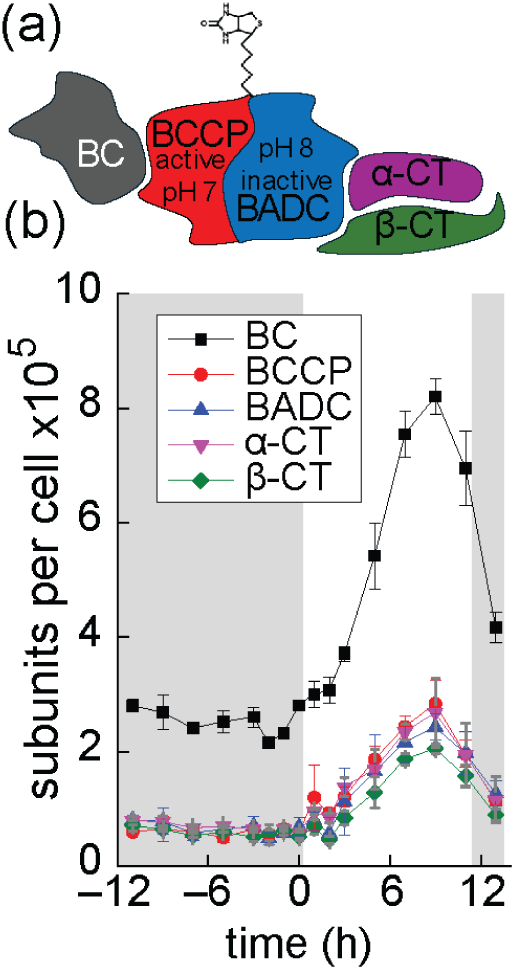
Copy number estimates of chloroplast ACCase. Top: Schematic showing a cartoon depicting the heteromeric ACCase complex. The abundance of each protein (in units of millions of molecules per cell) was estimated by quantitative proteomics using the proteomic ruler method and used to calculate the amount of each of the individual proteins (BC, BCCP, BADC, α-CT and β-CT) per cell over the cell cycle.

#### On the relationship between mRNA and protein levels along the cell cycle

We next sought to explore the relationship between transcript levels (determined by RNA-seq) and absolute protein abundances for all RNA/protein pairs along the diurnal cycle. For this purpose, we first divided all transcripts (from nuclear, plastid and mitochondrial genes) into 10 quantiles, based on their average expression estimates along the Chlamydomonas diurnal cycle (Supplemental Figure 3a) (52). Protein identification was skewed towards high-abundance transcripts, ∼70% of proteins were identified with 2 unique peptides originating from transcripts in the top decile of abundances, while only ∼10% proteins were captured from transcripts in the bottom decile (Supplemental Figure 3b). When taking into account variance of protein quantitation between replicates and completeness of the time course (Supplemental Figure 3b), the datasets were even more skewed towards higher abundance transcripts: ∼ 50% of proteins were still identified from genes corresponding to transcripts in the top 10% of the expression estimates (<50% average RSD, identified in every timepoint), while only 22 proteins (<1%) were found among the genes with expression estimates in the bottom 10% (average < 0.1 FPKM). Protein abundances were widely distributed in all transcript quantiles, but, in quantiles above an average of 2.5 FPKM, the abundance distributions shifted towards higher protein abundances (Supplemental Figure 3c). The other way around, proteins in the bottom decile of protein abundance based on our proteomics data, while still associated with a wide range of average transcript abundances, were generally associated with an increase in the distribution of transcript abundances along protein abundance quantiles (Supplemental Figure 3d).

We next used the part of the proteome dataset of 2266 proteins for which we obtained experimental data in every time point (av. RSD < 50%) and compared their protein abundance profiles to their transcript abundance profiles (30). To minimize potential biases related to not comparable methodologies for RNA and protein quantification and to address the differences in dynamic range between mRNA and protein abundances, we calculated mean centered and scaled values of mRNA and protein abundances for each time course (z-scores). We again used a hierarchical clustering approach leading to 16 clusters with distinct patterns of protein abundances and plotted them based on their peak along the diurnal cycle. Within each protein detailed look at the RNA profiles resulting in similar protein accumulation patterns. We identified, based on protein cluster size, 3-11 individual transcript profiles associated with each respective protein, using a secondary hierarchical clustering approach (Figure 7, Supplemental Table 4, Supplemental Figures 4-19). Again, the majority of proteins with complete experimental data in all time points showed an exact doubling in protein abundances during the day (∼90%, cluster 4-16, Figure 8), comparable to what has been observed in the larger dataset including incomplete time series (Figure 1d). This was most likely because the complete series was further biased towards more abundant proteins, removing some more transiently expressed, lower abundance proteins from the dataset that do not necessarily double in mass during the diurnal cycle, like for example PsbS (Figure 1c, Supplemental Figure 4b) or proteins involved in DNA replication like RNR, the DNA helicase and DNA polymerase subunits.

**Figure 8.**
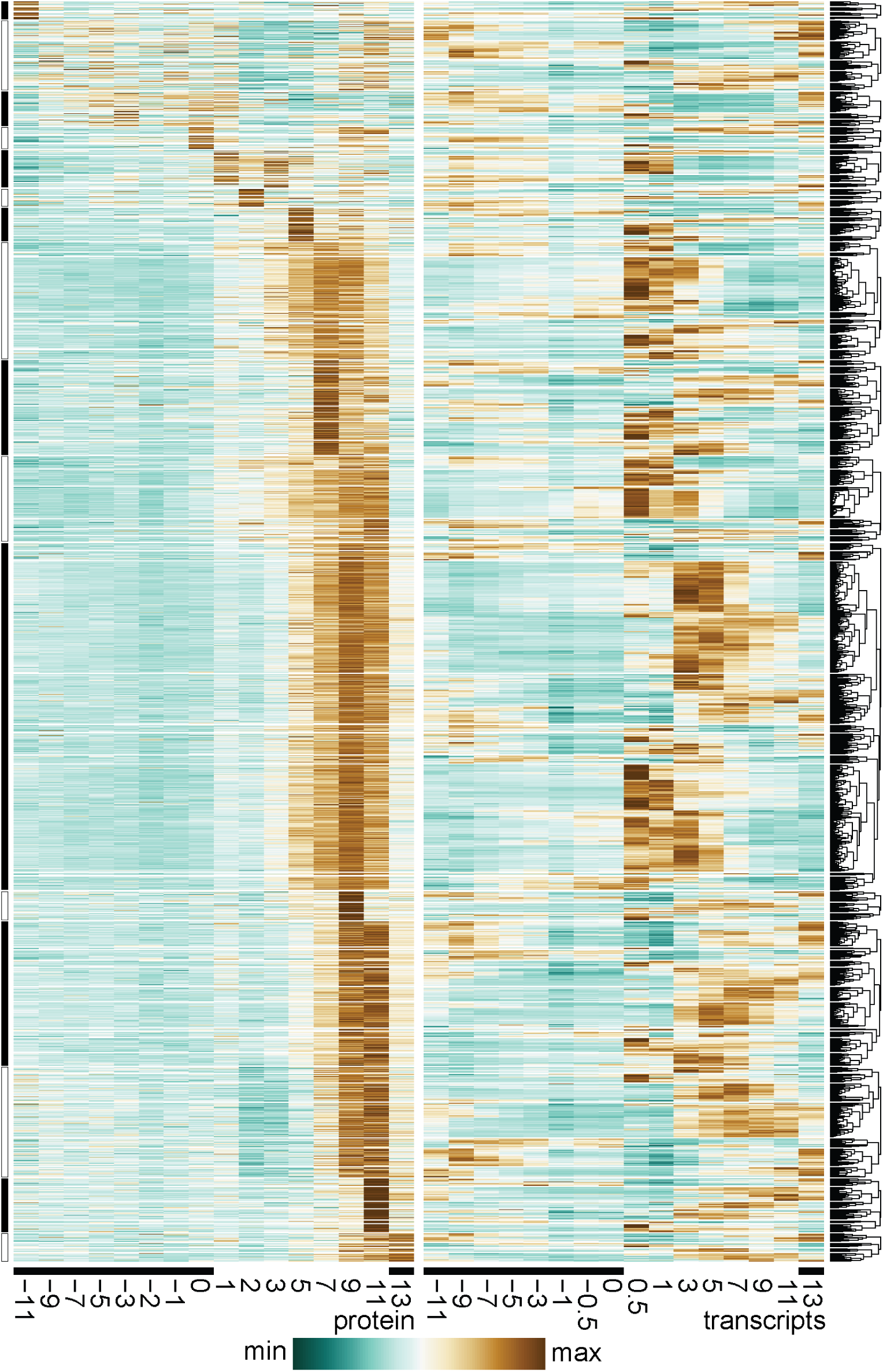
Identifying different transcript profiles amongst similar protein readouts. Heatmap representation of protein abundances for 2266 proteins (2 unique peptides, av RSD <50%, complete series) over one diurnal cycle and their corresponding transcripts. Proteins were clustered hierarchically and organized by peak abundance into 16 groups, each of which was again hierarchically clustered into 3-11 individual clusters based on the transcript abundance. Protein and transcript abundances for this figure were standardized (z-score).

As expected, transcript abundances had a greater dynamic range as compared to the protein abundances and showed more dramatic oscillation over the course of the dark light cycle, increasing contrast and allowing determination of the timing of induction more precisely. Sub-clustering for most protein clusters revealed a set of associated transcript profiles. Most transcript subclusters peaked in expression prior to protein accumulation, but with some variance in peak timing, or showing more transient or more sustained expression. For example, in protein cluster 11, which was the biggest cluster of the dataset containing 674 proteins accumulating over the day (peak at 9h light), transcript abundance profiles of four prominent subclusters (11.7 - 11.10, accounting for 354 of 674 proteins, ∼ 53 %) all peaked ∼ 3-5h in the light. While transcripts in subcluster 11.9 had a very sharp peak around 3/5h in the light, those in 11.7, starting also around 3h light, sustained transcript production until the end of the day. Transcripts in cluster 11.8 and 11.10 already increased 30 min into the day and were either sharply reduced after 5h of light (subcluster 11.8) or were sustained longer (11.10). We therefore conservatively classified the individual transcript subclusters into expression profiles that generally support the observed protein accumulation (green graph outlines, Supplemental Figure 4-19, green background, Supplementary table 4) or those that showed profiles that were less compatible (orange outlines in the figures and red numbers in the Supplemental Table 4).

We found that the vast majority of transcript profiles were compatible with the observed protein accumulation profiles, and that there is usually only one, sometimes two, transcriptional profiles that would require the utilization of other regulatory mechanism to achieve the observed protein production. This could be realized for example by using targeted protein degradation in parts of the diurnal cycle, or by employing translational mechanisms to increase translation initiation, efficiency, and fidelity rather than scheduling. In total we identified 306 transcript/protein pairs (out of 2266, ∼14%) in this category. In general, our study aligns well with work done in *Saccharomyces cerevisiae* which suggested that for ∼70% of the genes, the protein concentrations correlate well with the RNA concentrations (80). Despite our observed differences in abundance patterns between both biomolecules in a subset of cases, our results highlight the importance of regulation through mRNA abundance in biological systems.

## DISCUSSION

This work paints a quantitative picture of proteome dynamics over the cell cycle in Chlamydomonas cells and provides a workflow for generating and interpreting proteomics data using the TPA. TPA is compatible with data-dependent and data-independent acquisition MS workflows, using spectral intensities to calibrate label-free, large-scale proteomics datasets to the total cellular protein amount to obtain absolute protein abundances without the use of external standards (25, 81). As such, it does not require prior hypotheses, or expensive, isotopically-defined protein/peptide standards, and can even be applied on existing datasets when the cellular protein content of a sample is known. With this workflow, we quantitatively captured copy number estimates of dozens of subunits of chloroplast multi-subunit protein complexes whose abundances can range over two orders of magnitude; both in the thylakoid membrane and the stroma. TPA relies on depth of coverage so that there is adequate representation of peptides throughout the dataset for each protein of interest, especially in data-dependent workflows. The ability to predict peptide observability is somewhat limited but has several important implications, especially when it comes to absolute protein quantification with mass spectrometry. Experimentally observed intensities of peptides that were preselected based on favorable physicochemical properties do not consistently correlate well with predicted peptide intensities (Pearson correlation coefficient of 0.62, (82)), which means ideally the peptide chosen for quantitation has already been observed in prior experiments and found to be accumulating well before selecting its use for quantitation. TPA instead uses averages between multiple peptides, either random or a select subset of peptides (for example most abundant). Certain types of proteins are poorly represented in data-dependent proteomic datasets in general, such as small proteins (few discoverable peptides), highly hydrophobic multi-spanning transmembrane proteins or proteins with uneven Arg and Lys distribution and low abundance proteins. This means that there is either a lower confidence in the absolute abundance of these proteins as determined with TPA, or the amount of protein is underestimated by the method, and together with the bias of data-dependent acquisition to favor more abundant proteins, studies utilizing TPA will find better results for abundant multiprotein complexes than for small, individual, low-abundant proteins, which is also what we observed (Figure 1), results from less reliable coverage and accuracy.

For abundant complexes however, TPA appears to be quite accurate. We tested the validity of the obtained abundance estimates by comparing them to other absolute quantifiable markers, e.g. using the total Cu content of cells. Under the growth conditions used in this work, two proteins, highly abundant plastocyanin and less abundant Cyt oxidase, determine the Cu content of the cells (37), which can be independently and accurately quantified against a Cu standard by inductively coupled plasma-mass spectrometry (ICP-MS/MS). The match between the protein and the metal measurements offers confidence in the estimated derived from TPA, both in terms of their absolute amounts but also in their repeatability between time points, especially in the dark where there is little metabolic activity after cell division, and in their accumulation dynamics during the day. Furthermore, the copy number of plastocyanin measured in this work is in the same range as previous estimates (18) using isotopically-defined standards. The same is true for the observed correlation between Chl content, that can be quantified accurately spectroscopically, and the abundance of Chl binding proteins, again, speaking for the accuracy in protein quantification using the TPA. The ability to accurately quantify photosynthetic pigments and trace metal cofactors involved in photosynthesis might be a unique advantage for absolute protein abundance determination in proteomics experiments in phototrophic organisms. Photosynthetic proteins are highly abundant and likely to be consistently recovered in these experiments, the photosynthetic protein complexes are highly conserved between phototrophs, and with the advent of CryoEM approaches, many structural studies have detailed the stoichiometries and composition of photosynthetic complexes (55, 83). Furthermore, photosynthetic complexes of oxygenic phototrophs are multi-subunit complexes that are more resilient to a miss in coverage in individual time points/samples in data-dependent workflows.

For the photosynthetic apparatus, we estimate the following copy numbers per cell and stoichiometries (Table1): Rubisco, PSII, PSI, Cyt *b*_6_*f*, ATP synthase at 5:2:2:1:1 and 0.98 × 10^6^ molecules per cell and for the electron transfer chain we observe ratios of 3:100:1:3.2 for PSII, PQ, Cyt *b*_6_*f*, plastocyanin and PSI. We noted that the mobile electron carrier PQ and the oxygen scavenger α-tocopherol are found at relatively high excess relative to PSII (31:1 PQ:PSII and 13:1 α-tocopherol:PSII). The latter is not that surprising given that cells need to keep generating α-tocopherol as it is required to scavenge reactive oxygen species, but irreversibly consumed after being oxidized during singlet oxygen scavenging. The number of PQ molecules per PSII has previously been estimated to be ∼15:1 but is dependent on the growth regime (84, 85). The excess PQ observed by us and others may reflect additional PQ outside the thylakoid membranes in addition to photoactive PQ pool. It was proposed that this additional PQ makes up between 30 and 50% of the total PQ content and is involved in quenching reactive oxygen species (85, 86).

An interesting finding from a similar chemical analysis of quinones and tocopherols is the 1:1 stoichiometric occurrence of γ-tocopherol and the PSII core complex, suggesting that γ-tocopherol could be an intrinsic component of PSII or at least associated with one of the Psb polypeptides (83).

PSI and PSII are stoichiometrically more abundant than are the Cyt *b*_6_*f* and ATP synthase complexes throughout the cell cycle. Given their lower turnover relative to the photosystems, this might suggest that these proton-pumping complexes would be rate limiting for energy transduction. Nevertheless, the stoichiometries measured in the laboratory may reflect adaptation of Chlamydomonas to the low photon flux density of its natural soil habitat. In this context, one might expect that photosystem core complexes and antenna abundances might decrease under high light acclimation (87).

Besides estimating the number of each complex per cell we could also determine the subunit composition of multi-protein enzymes or in other words the stoichiometry of each polypeptide in a macromolecular complex. For Rubisco, we recapitulated the well-known 1:1 stoichiometry, for CF_1_, the 3:3:1 stoichiometry for the alpha, beta and gamma subunits and for the hexameric FtsH protease, the 1:1 stoichiometry of the A and B type subunits. On the other hand, for the ClpP proteases we noted the heptameric upper ring was recovered with a stoichiometry different from that noted in the structural work. This can happen if one or more subunits are under-represented in the dataset. Nevertheless, we ruled out that possibility because the total number of top and bottom ring polypeptides is well matched (Figure 4). It is rather more likely that there is compositional heterogeneity in the chloroplast Clp protease. This experimental design also reveals heterogeneity during the course of the day as we noted for the Lhcbm3 and Lhcbm4 polypeptides whose increase is coincident with the increase of Chl *b*, speaking to changes in the organization of the PSII accessory antenna. It should be possible to isolate PSII-LHCII super-complexes over the course of the day to visualize these changes by structural analysis. The same is true for the abundance of Lhca6, which is induced more than 4.4fold during the day as compared to the expected, average ∼2fold increase observed for all other photosynthetic complexes and antenna proteins (Supplemental Table 3).

TPA is a high throughput genome-wide approach that does not require preparation of individual reagents (e.g. antibodies) for proteins of interest nor biochemical purifications. With copy number estimates for entire pathways of interest, quantification by TPA generates datasets that can be exploited for building metabolic models in which values for active site concentrations can be incorporated in relation to metabolite concentrations. Certainly, individual subunits of complexes may be over- or under-represented by TPA as compared to classical labeling approaches or structure determination. Nevertheless, a key advantage of the TPA is that the data are generated on total cell extracts (see Methods) without additional fractionation and hence less subject to selective loss of peptides from the pool. We note also that any *in vivo* subunit heterogeneity of multi-protein complexes can be obscured by application of methods that emphasize homogeneity such as biochemical preparations or crystallography or by application of methods that rely on signal averaging as in cryo-EM. With respect to copy number estimates of multi-subunit complexes, we suggest that the TPA is rather robust because it is the result of quantification of multiple individual subunits.

Finally, we use the quantitative proteomics data set and the previous RNA-Seq datasets, both with high temporal resolution to examine the relationship between cellular mRNA and protein abundances at a genome-wide level. We identify several distinct patterns. The abundance of proteins that contribute to biomass track with the increase in cell size and total protein during G1, like the major photosystems, and proteins involved in carbon assimilation and metabolism like Rubisco and carbonic anhydrases. Within this group the mRNAs show temporally phased peaks of expression (30, 31) with proteins accumulating linearly during the G1 phase reflecting the long half-lives of these proteins (e.g. histones, proteins of the photosynthetic apparatus, enzymes of Chl biosynthesis) and the necessity of maintaining copy numbers post-division. With approximately 10^5^ mRNAs per cell, perhaps 7 ribosomes per polysome and approximately 10^6^ 80S ribosomes, the temporal sharp rise and decay of groups of mRNAs for particular macromolecular processes (Chl biosynthesis, DNA replication, cilia biogenesis) that occur at specific points in the cell cycle are critical for ensuring the synthesis of the corresponding proteins for executing the cell cycle program. Proteins that catalyze specific cell cycle events such as DNA synthesis during the S phase or during mitosis show a precisely timed peak of accumulation that typically follows a similar peak in accumulation of the corresponding mRNA (e.g. ribonucleotide reductase, DNA polymerase) (30). For these situations, changes in mRNA and protein half-life are likely key determinants of RNA and protein abundance. We also identify several mRNAs whose abundance is stable over the course of the day, but whose corresponding polypeptides exhibit dynamic changes in abundance (Supplemental Table 2). Multi-layered control points, such as transcription of RNA and post-translational degradation of the protein can affect more nimble responses to stimuli and yet tighter control points.

## EXPERIMENTAL PROCEDURES

### Culture conditions

*Chlamydomonas reinhardtii* strain CC5390 was grown in flat panel photo-bioreactors, exactly as described in (52). Replicate samples were always taken from independent photo-bioreactors for various measurements as described below.

### Protein detection by LC-MS/MS

We collected 10^8^ cells by centrifugation at 1,450 *g* at 4°C for 4 min. The cell pellet was resuspended in 200 µl 10 mM Na-phosphate, pH 7.0 and flash frozen in liquid nitrogen. Protein concentration was determined by BCA assay (Thermo Scientific, MA, USA). Urea and dithiothreitol were added to all samples at a final concentration of 8 M and 10 mM, respectively before incubation at 60°C for 30 min with constant shaking (800 rpm). All samples were then diluted 8-fold with 100 mM NH_4_HCO_3_ and 1 mM CaCl_2_ and digested with sequencing-grade modified porcine trypsin (Promega, WI, USA) provided at a 1:50 [w/w] trypsin-to protein ratio for 3 h at 37°C. Digested samples were desalted using a 4-probe positive pressure Gilson GX-274 ASPEC™ system (Gilson Inc., WI, USA) with Discovery C18 100 mg/mL solid phase extraction tubes (Supelco, MO, USA) as follows: columns were pre-conditioned with 3 ml methanol, followed by 2 ml 0.1%trifluoroacetic acid (TFA) in water. Samples were then loaded onto columns, followed by a wash using 4 ml 95:5 water: acetonitrile (ACN) 0.1% TFA. Samples were eluted with 1 ml 20:80 water:ACN 0.1% TFA, and concentrated to a final volume of ∼100 µl in a Speed Vac. After determination of peptide concentration by BCA assay, samples were diluted to 0.25 µg/µl with nanopore water for LC-MS/MS analysis (LC part: LC column of fused silica [360 µm x 70 cm] handpacked with Phenomenex Jupiter derivatized silica beads of 3 µm pore size (Phenomenex, CA, USA); HPLC part: HPLC NanoAcquity UPLC system (Waters, MA, USA); MS part: Q Exactive Plus mass spectrometer (Thermo Fisher, MA, USA)). Samples were loaded onto LC columns with 0.05% formic acid in water and eluted in 0.05% formic acid in ACN over 100 min. Twelve high resolution (17.5K nominal resolution) data-dependent MS/MS scans were recorded in centroid mode for each survey MS scan (35K nominal resolution) using normalized collision energy of 30, isolation width of 2.0 m/z, and rolling exclusion window lasting 30 seconds before previously fragmented signals are eligible for re-analysis. Unassigned charge and singly charged precursor ions were ignored.

### MS/MS data search

The MS/MS spectra from all LC-MS/MS datasets were converted to ASCII text (.dta format) using MSConvert (http://proteowizard.sourceforge.net/tools/msconvert.html) which precisely assigns the charge and parent mass values to an MS/MS spectrum as well as converting them to centroid. The data files were then interrogated via target-decoy approach (88) using MSGFPlus (89) using a +/− 20 ppm parent mass tolerance, partial tryptic enzyme settings, and a variable posttranslational modification of oxidized Methionine. All MS/MS search results for each dataset were collated into tab separated ASCII text files listing the best scoring identification for each spectrum.

### Data analysis

MS/MS search results for each biological replicate were collated to a tab delimited text file using in-house program MAGE Extractor. These results were then imported into a SQL Server (Microsoft, Redmond, WA) database and filtered to 1% FDR by adjusting the Q-Value provided by MSGF+. The in-house program MASIC (https://omics.pnl.gov/software/masic, 53) was run on each dataset, which provides ion statistics and general information extracted from the instrument binary rawfile. Specifically, both the maximum observed MS ion count for each MS/MS spectrum’s precursor signal (termed PeakMaxIntensity) and a selected ion chromatogram (SIC, termed StatMomentsArea) are provided for subsequent peptide quantitation. The results from all datasets’ MASIC output were collated using the in-house program MAGE File Processor, with a tab delimited text file listing all relevant data as the result. These results were also imported into SQL Server and connected to the filtered MSGF+ results via Dataset_ID (internal to PNNL’s data management system), and relevant MS/MS scan number. Unique peptide sequences (with post-translational modifications counted as separate sequences) were grouped per dataset with observation counts and maximum PeakMaxIntensity values provided. This data was then pivoted to provide a crosstab of peptides, with protein information carried through the query. The crosstab was then imported into InfernoRDN (an implementation of the R statistical package, https://omics.pnl.gov/software/infernordn), Log_2_ transformed and mean central tendency normalized (boxplot alignment). The normalized peptide abundance values were then exported into Excel (Microsoft), anti-logged, and proteins grouped with values summed using the Pivot Table function. Most further analysis were carried out in R, including the determination of euclidean distances (stats package) and hierarchical clustering (stats package, using the ward.D2 algorithm). Graphing was performed in R (ggplot2, pheatmap packages) and further modified in Adobe Illustrator. The mass spectrometry proteomics data have been deposited in the ProteomeXchange Consortium (www.proteomexchange.org/) under accession no. PXD020726 and will be made publicly available upon publication.

### Elemental content measurements by ICP-MS/MS

We collected 5×10^7^ cells by centrifugation at 3,500 *g* for 3 min and washed twice with 1 mM Na_2_-EDTA, pH 8.0, to remove metals associated with the cell surface and twice with Milli-Q water. After removing the remaining water by brief centrifugation, cell pellets were digested with 70% nitric acid at room temperature overnight and at 65 °C for 2 h. Digested samples were diluted with Milli-Q water to a final nitric acid concentration of 2% (v/v). To measure metal content of culture medium, aliquots of the medium were treated with nitric acid and diluted with Milli-Q water to reach a final concentration of 2% nitric acid (v/v). Elemental analysis was measured by inductively coupled plasma MS on an Agilent 8800 Triple Quadrupole ICP-MS instrument using three standards for calibration (an environmental calibration standard (Agilent 5183–4688), phosphorus standard (Inorganic Ventures CGP1), and sulfur standard (Inorganic Ventures CGS1)) and two internal standards (^89^Y and ^45^Sc (Inorganic Ventures MSY-100PPM and MSSC-100PPM, respectively)). Elements were determined in MS/MS mode and measured in a collision reaction cell using helium for the measurement of ^63^Cu. Each sample was measured in four technical replicates, and variation between technical replicates did not exceed 5%.

### Plastoquinone-9 and tocopherol analyses

PQ and tocopherols were quantified in the same cell extracts via reverse-phase chromatography as previously described (30, 90).

## ACKNOWLEDGMENTS

This work was supported by a grant to SSM from the National Institutes of Health (GM42143) and by National Science Foundation Grant MCB-2216747 to GJB. Proteomics work was performed at the Environmental Molecular Science Laboratory, a Department of Energy (DOE) Office of Science User Facility sponsored by the Office of Biological and Environmental Research (BER) and located at Pacific Northwest National Laboratory (PNNL), proposal ID 49840. Pacific Northwest National Lab is operated by Battelle for the DOE under Contract DE-AC05-76RL01830.

## SHORT LEGENDS FOR SUPPORTING INFORMATION

**Supplemental Table 1.** Protein quantification data. Relative protein abundance data shown as either spectral count (SC) or MASIC based protein abundance estimates.

**Supplemental Table 2.** Organized and labeled information on protein clusters and corresponding RNA abundance data.

**Supplemental Table 3.** Protein copy number estimates per cell. Shown are GRAVY numbers and calculations to estimate protein copy numbers /cell for selected proteins.

