## Supplemental Figures and Legends for "Estimation of chloroplast macromolecular complex copy numbers and subunit stoichiometries during the *Chlamydomonas reinhardtii* cell cycle"

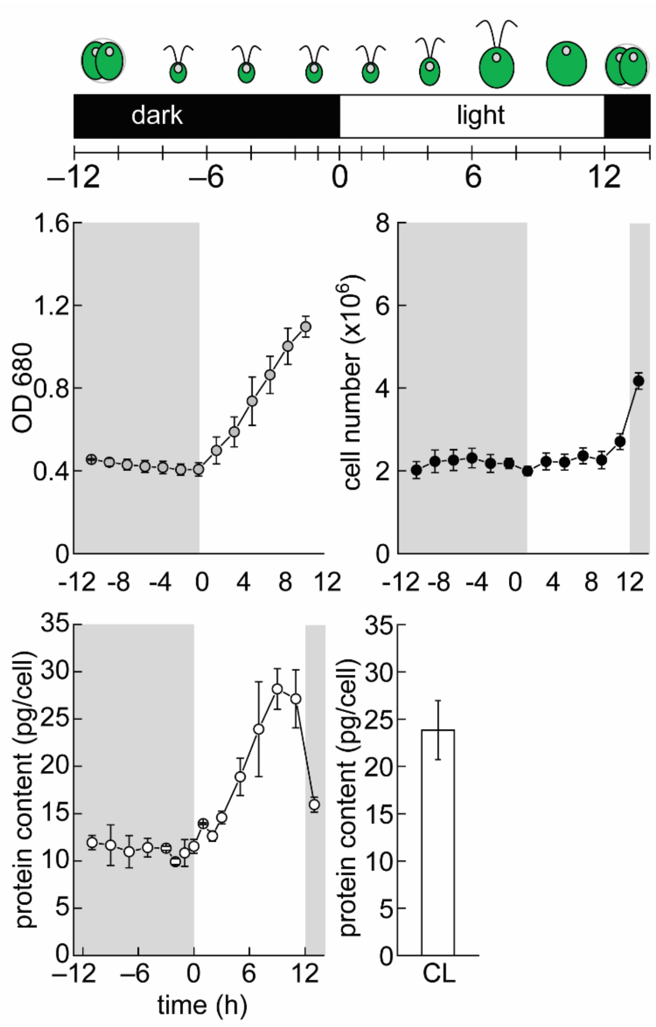


**Supplemental Figure 1.** Measurements of optical density (OD680), cell numbers using a hemocytometer and protein abundances using BCA assays. Shown are averages and StDev of samples from three independent experiments/photobioreactors.

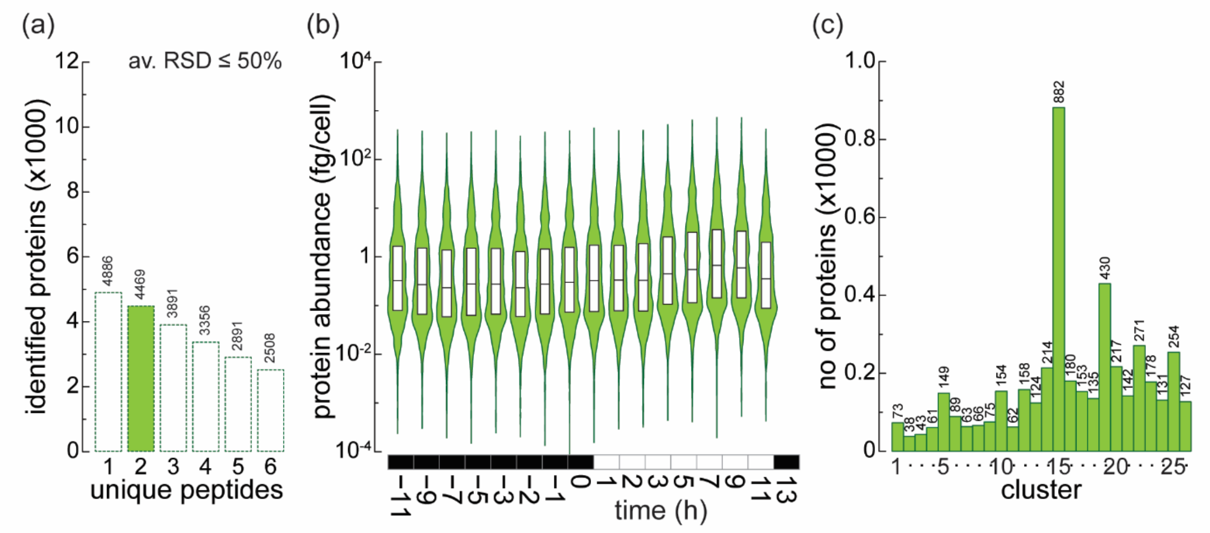


**Supplemental Figure 2.** (a) Number of proteins identified (av. RSD < 50%) in the dataset dependent on the number of required unique peptides for each protein. The used cutoff is highlighted by green fill. (b) Distribution of protein abundances in each timepoint are shown with violin plots, median, 25% and 75% quantile are shown with box overlays. (c) Overview of the hierarchical clustering

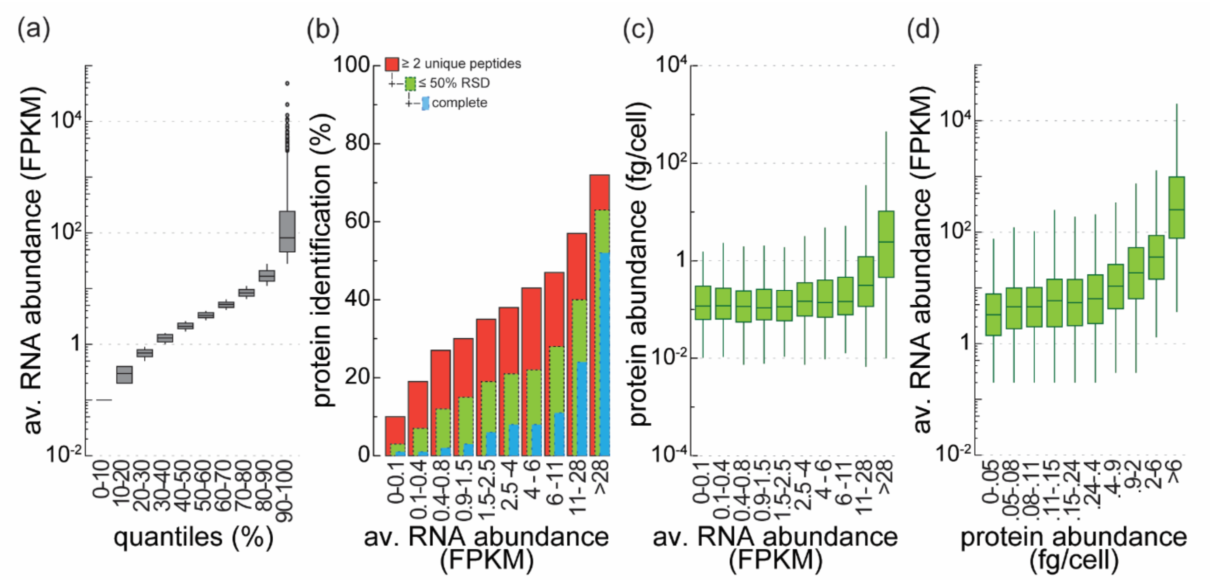


**Supplemental Figure 3:** (a) Range of average transcript abundances (FPKM) in 10 quantiles (10% each) in the diurnal cycle (b) Fraction of proteins identified in the proteomics experiment based on their corresponding transcript abundance. Red fill for 2 unique peptides and above, green fill for the fraction of proteins quantified with an av. RSD <50%, blue fill for those proteins captured at every time point (c) Protein abundance (fg/cell) associated with different transcript abundance quantiles (d) Transcript abundance associated with different protein abundance quantiles

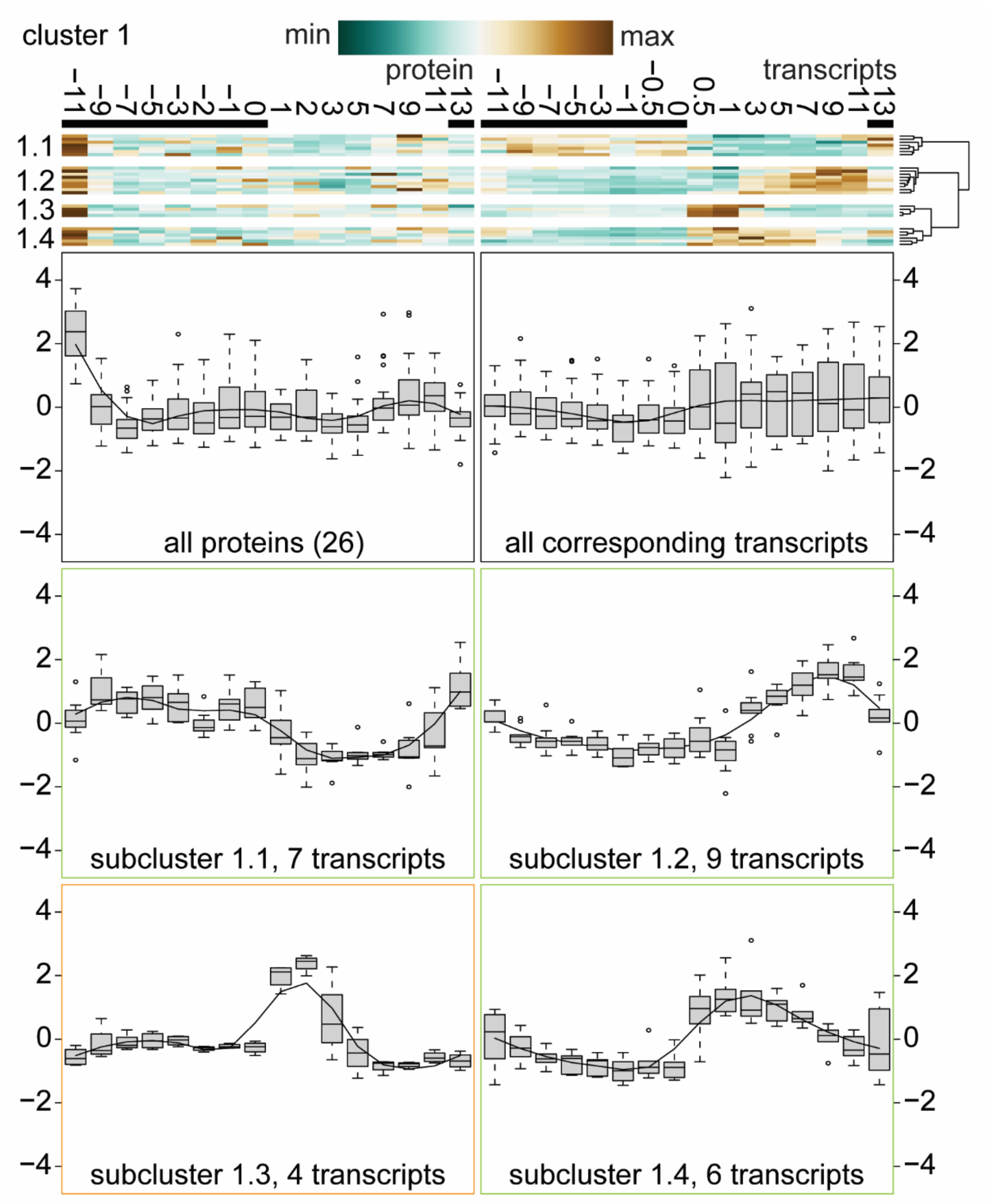


**Supplemental Figure 4:** Cluster 1 of 16, protein abundances peaking at -11 h early in the night and associated transcript profiles. Hierarchical clustering was performed first on protein abundances of proteins identified in every time point (av. RSD <50%) and then organized by their corresponding transcript profiles into 4 subclusters.

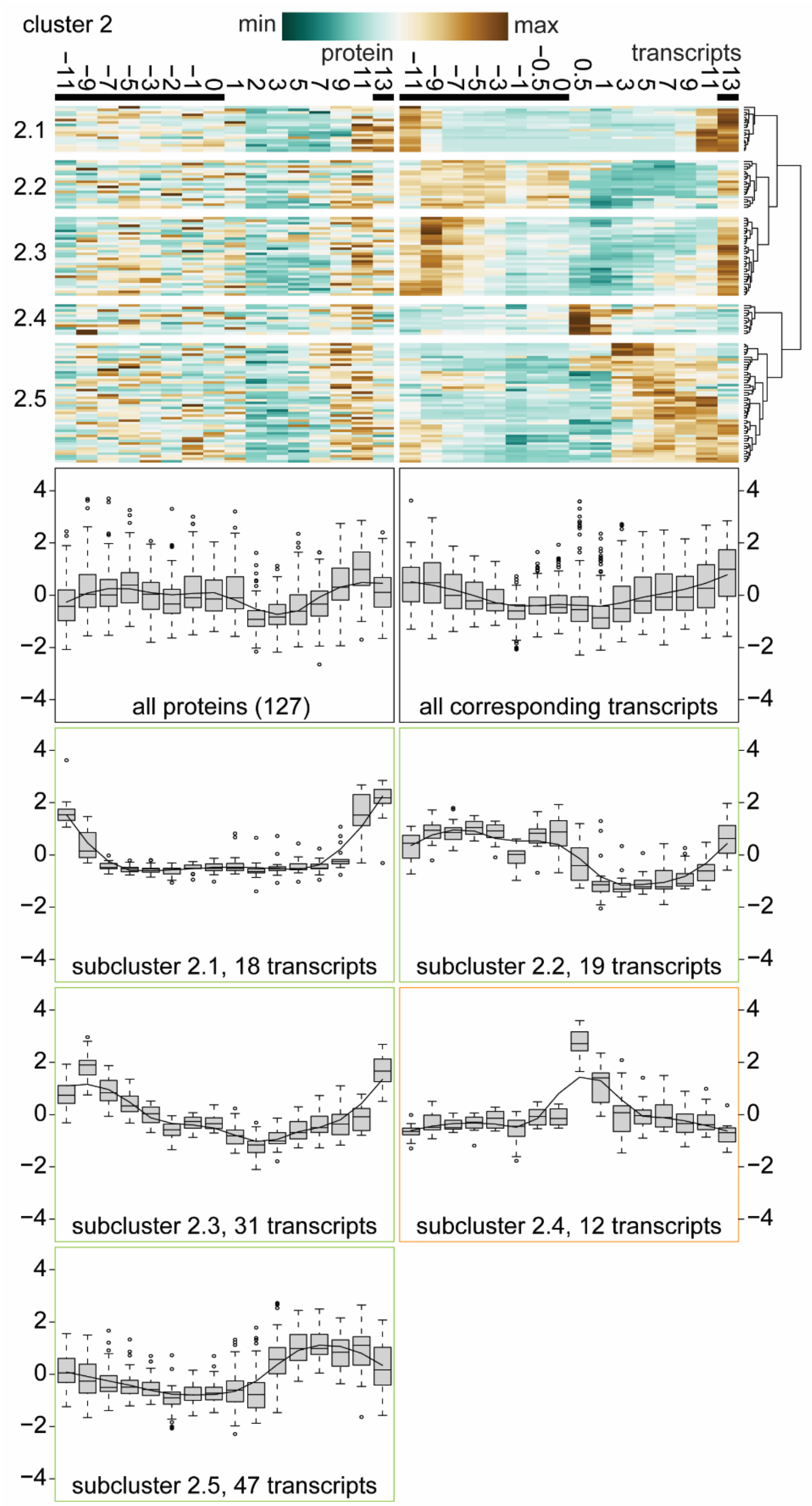


**Supplemental Figure 5:** Cluster 2 of 16, protein abundances reduced during the day, induced early in the night or very late in the day, and associated transcript profiles. Hierarchical clustering was performed first on protein abundances of proteins identified in every time point (av. RSD <50%) and then organized by their corresponding transcript profiles into 5 subclusters.

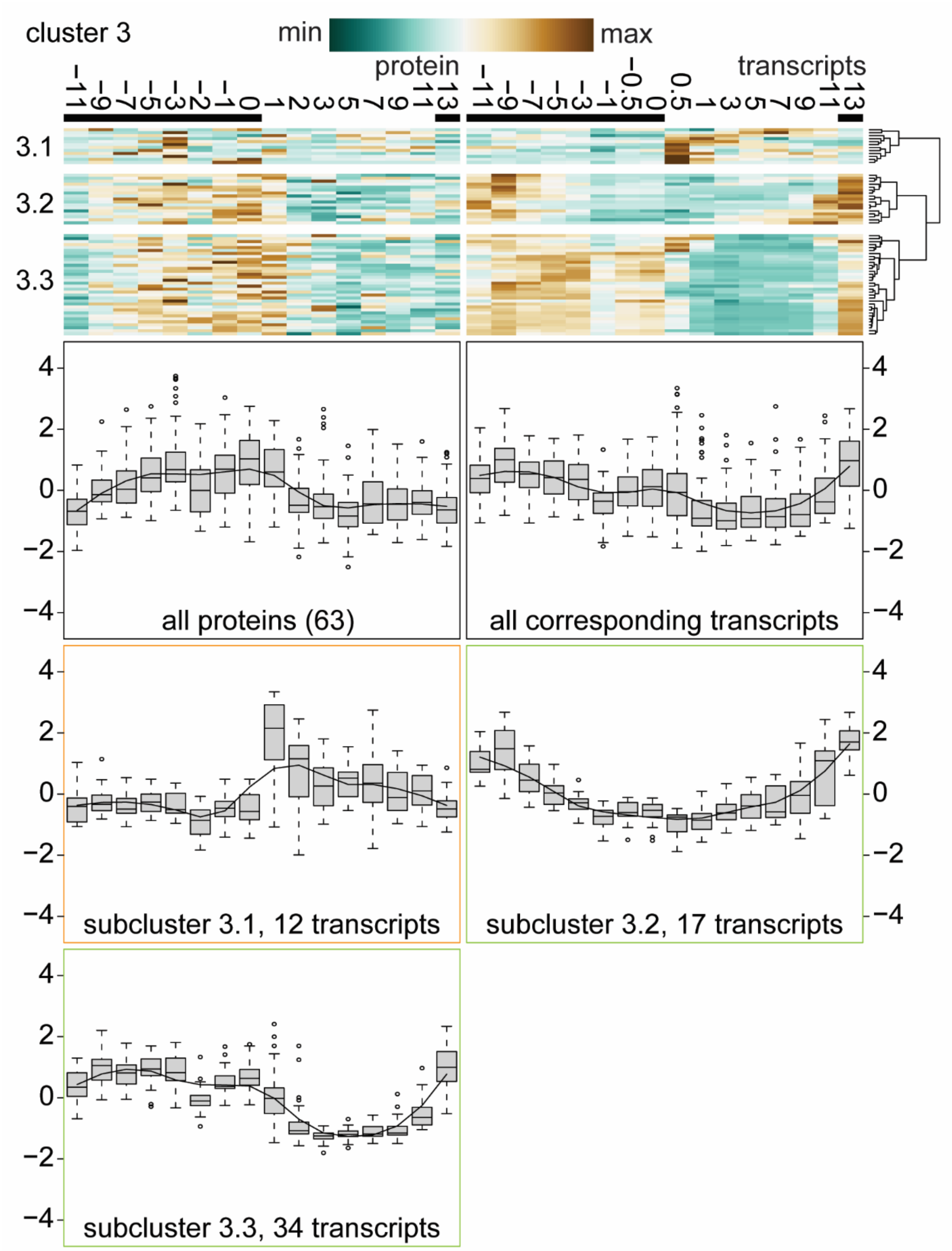


**Supplemental Figure 6:** Cluster 3 of 16, protein abundances reduced during the day, induction later in the night, and associated transcript profiles. Hierarchical clustering was performed first on protein abundances of proteins identified in every time point (av. RSD <50%) and then organized by their corresponding transcript profiles into 3 subclusters.

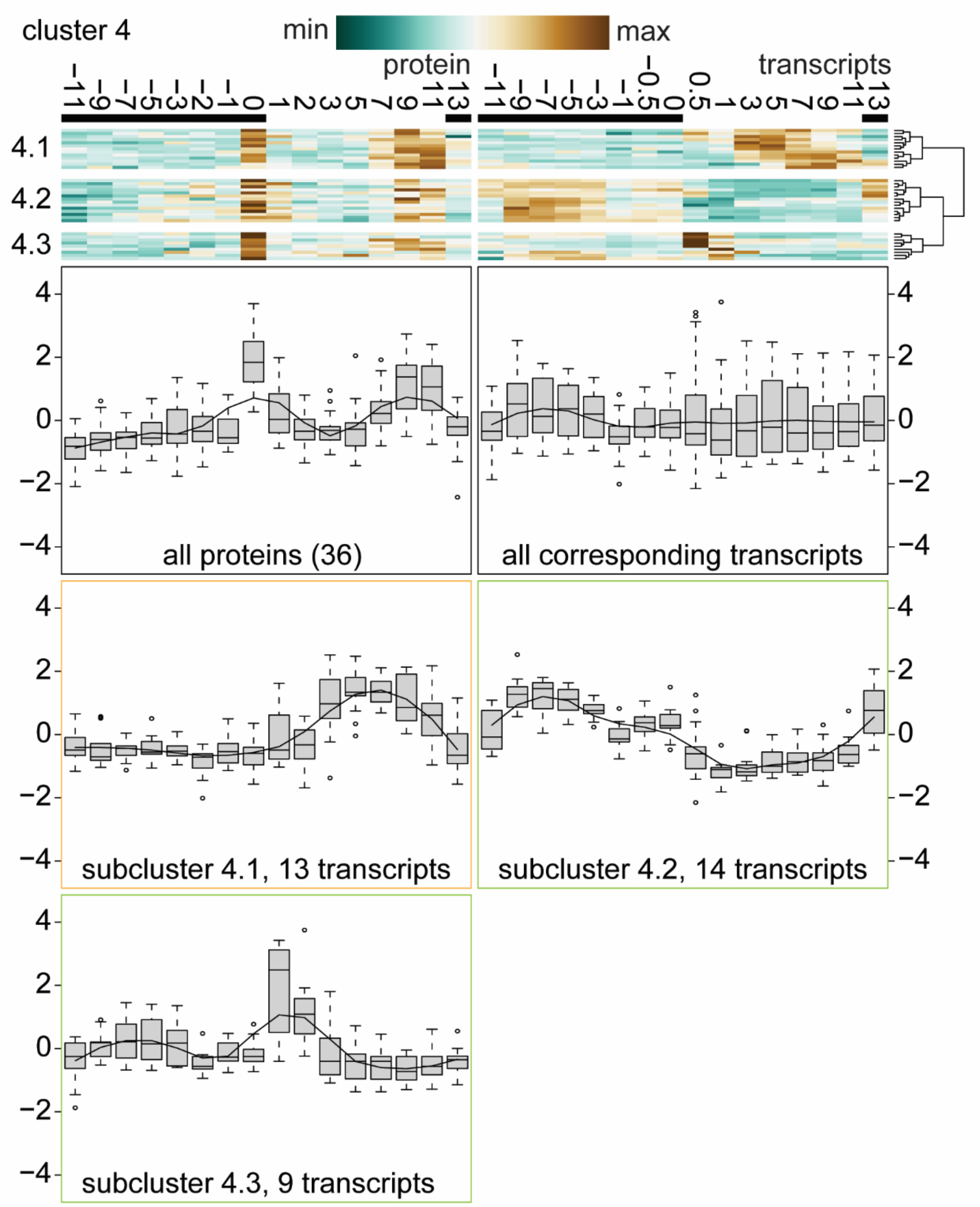


**Supplemental Figure 7:** Cluster 4 of 16, protein abundances transiently peaking at the onset of the day, secondary peak towards the end of the day, and associated transcript profiles. Hierarchical clustering was performed first on protein abundances of proteins identified in every time point (av. RSD <50%) and then organized by their corresponding transcript profiles into 3 subclusters.

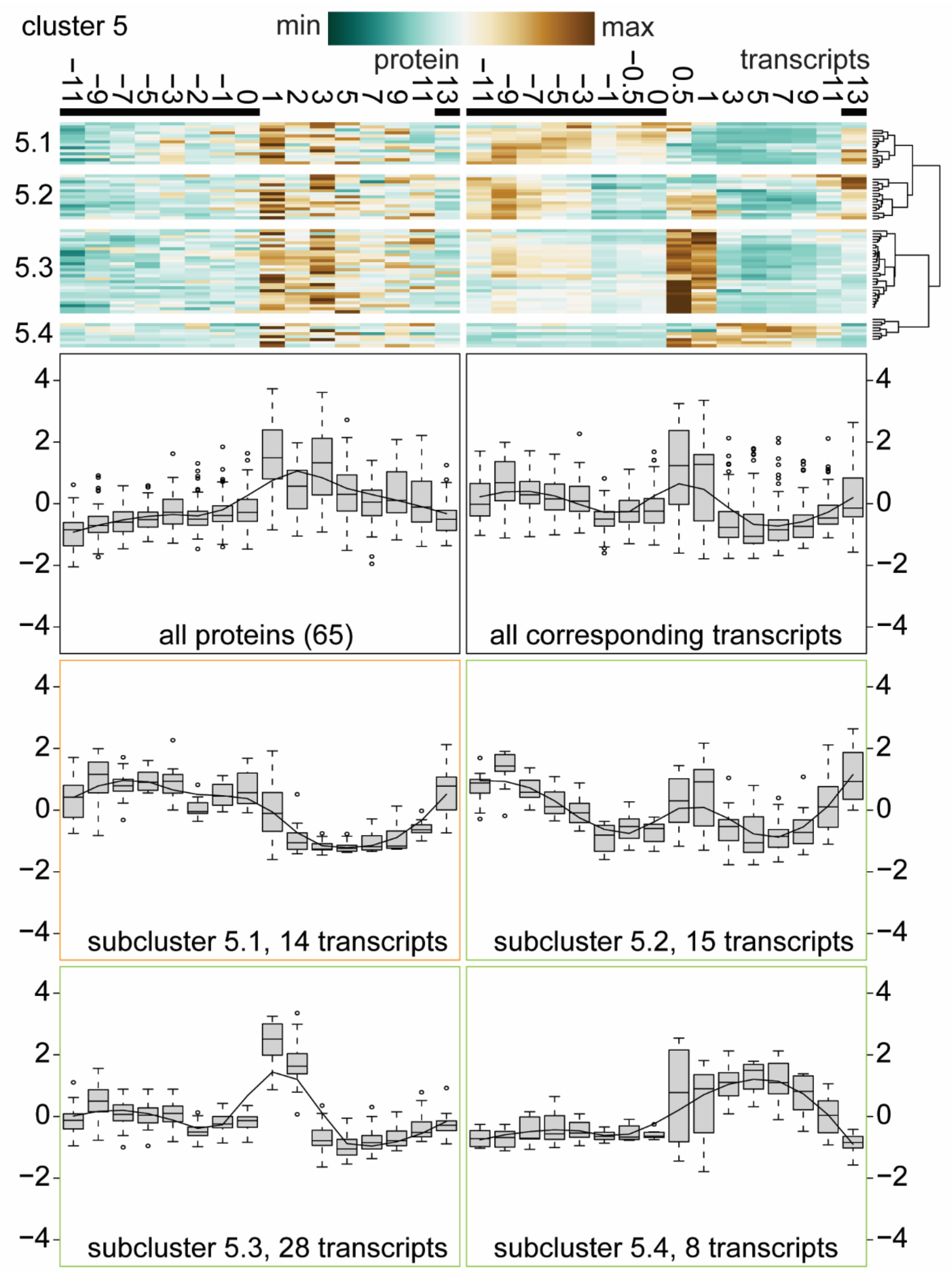


**Supplemental Figure 8:** Cluster 5 of 16, protein abundances peaking 1h in the day and associated transcript profiles. Hierarchical clustering was performed first on protein abundances of proteins identified in every time point (av. RSD <50%) and then organized by their corresponding transcript profiles into 4 subclusters.

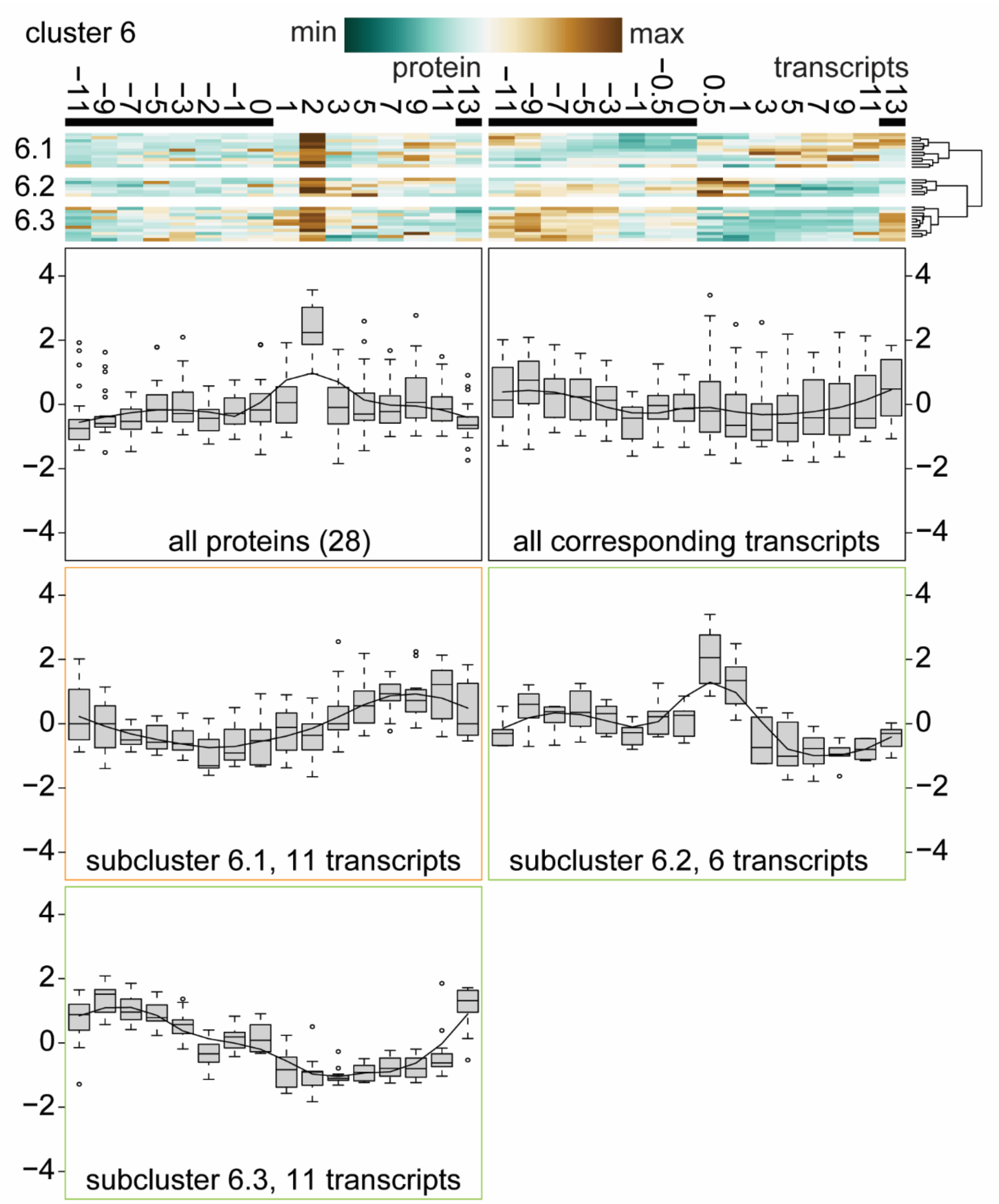


**Supplemental Figure 9:** Cluster 6 of 16, protein abundances transiently peaking 2h in the day and associated transcript profiles. Hierarchical clustering was performed first on protein abundances of proteins identified in every time point (av. RSD <50%) and then organized by their corresponding transcript profiles into 3 subclusters.

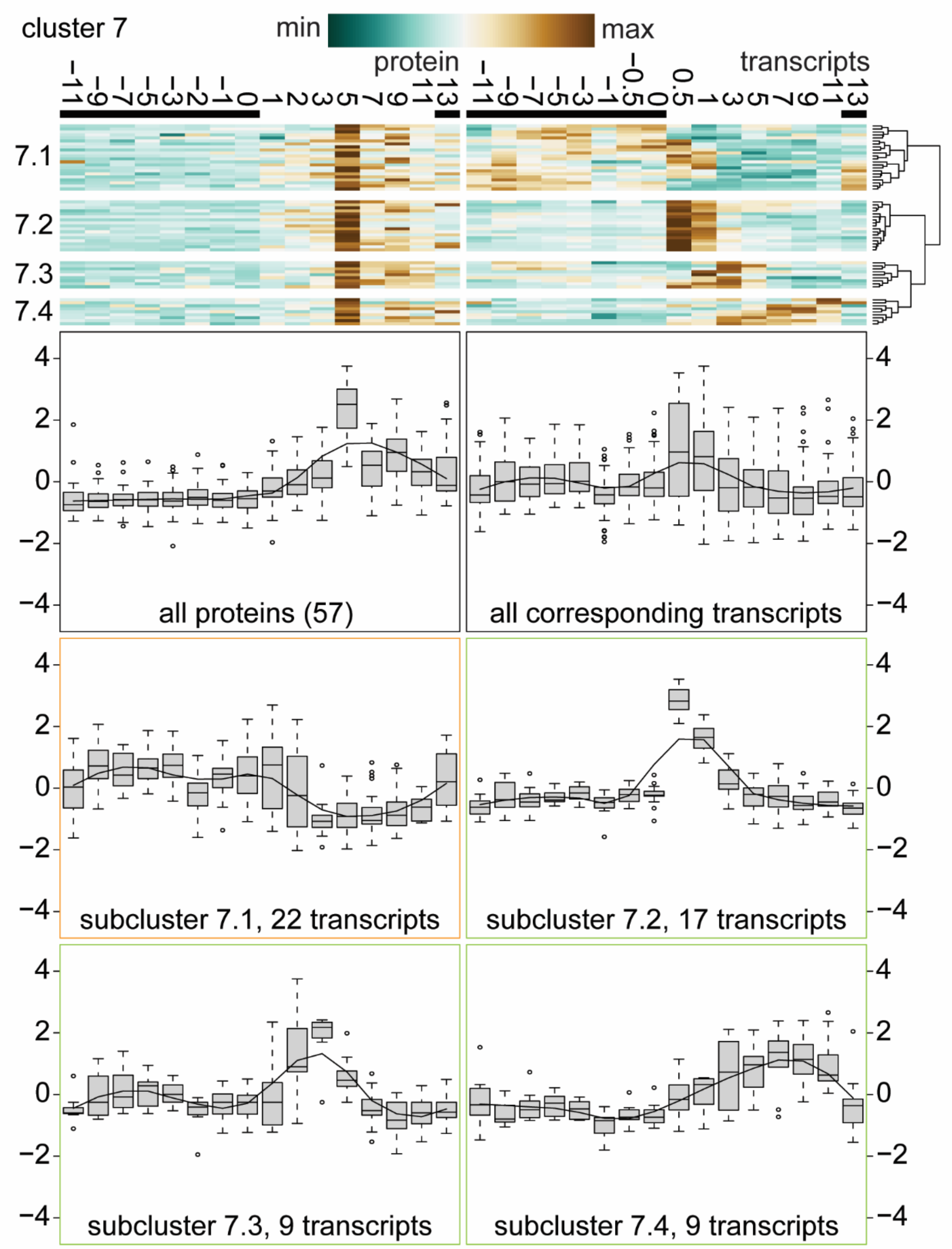


**Supplemental Figure 10:** Cluster 7 of 16, protein abundances transiently peaking 5h in the day and associated transcript profiles. Hierarchical clustering was performed first on protein abundances of proteins identified in every time point (av. RSD <50%) and then organized by their corresponding transcript profiles into 4 subclusters.

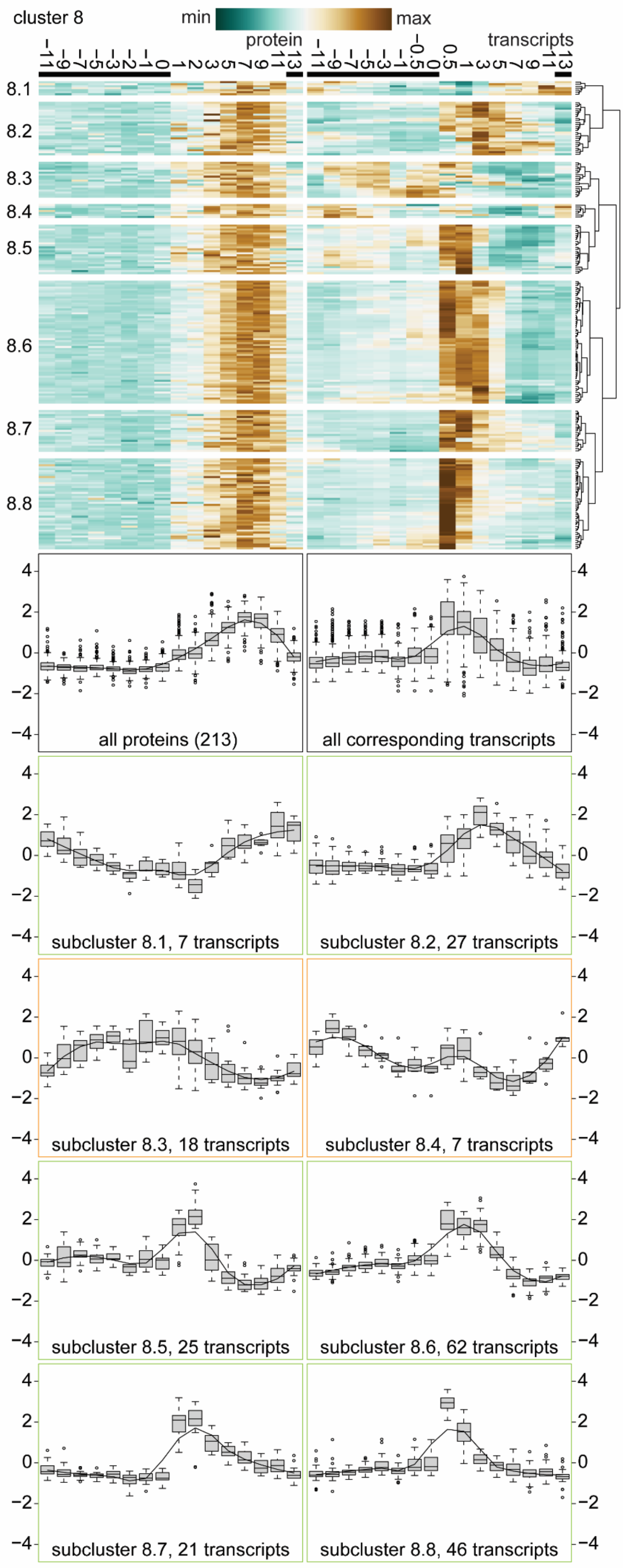


**Supplemental Figure 11:** Cluster 8 of 16, protein abundances peaking 7h in the day, earlier induction, and associated transcript profiles. Hierarchical clustering was performed first on protein abundances of proteins identified in every time point (av. RSD <50%) and then organized by their corresponding transcript profiles into 8 subclusters.

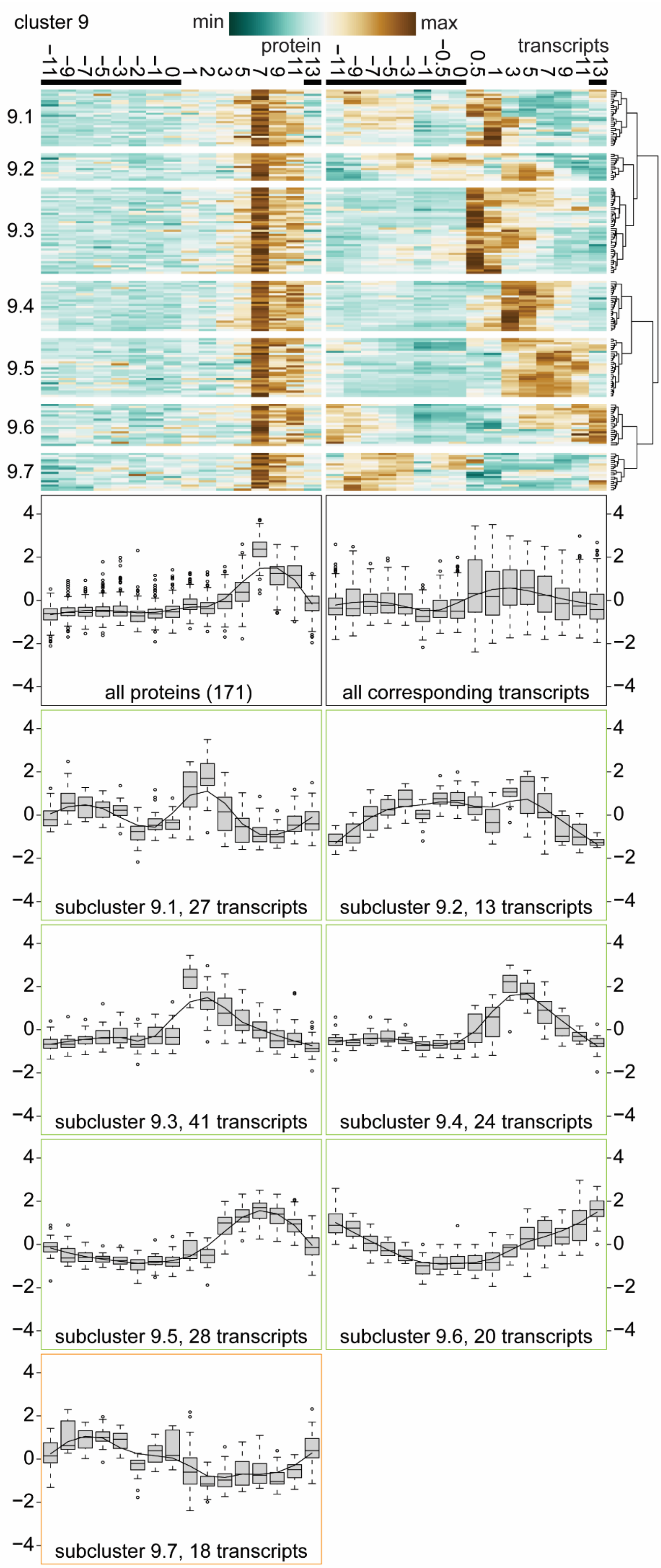


**Supplemental Figure 12:** Cluster 9 of 16, protein abundances peaking 7h in the day and associated transcript profiles. Hierarchical clustering was performed first on protein abundances of proteins identified in every time point (av. RSD <50%) and then organized by their corresponding transcript profiles into 7 subclusters.

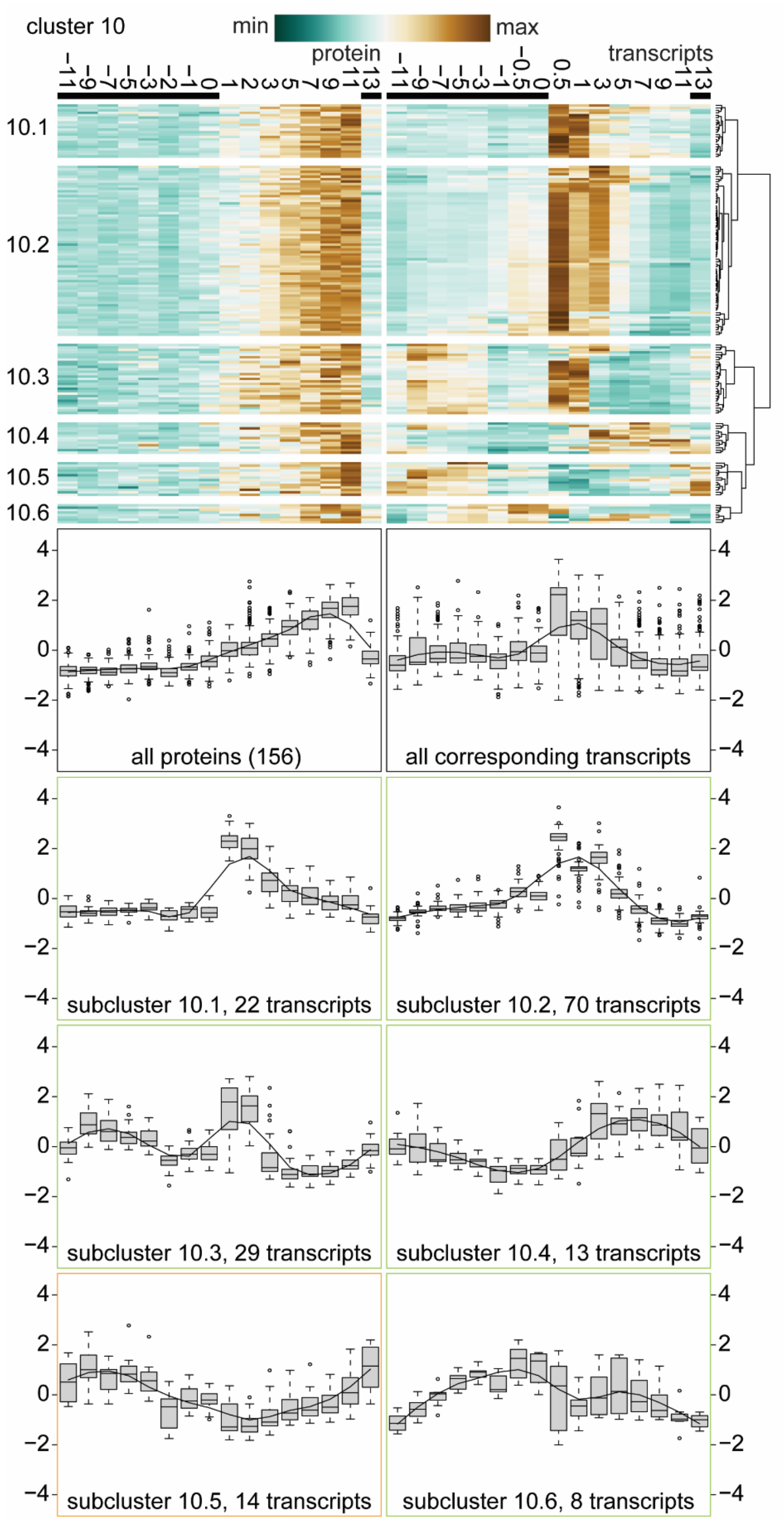


**Supplemental Figure 13:** Cluster 10 of 16, protein abundances peaking towards the end of the day and associated transcript profiles. Hierarchical clustering was performed first on protein abundances of proteins identified in every time point (av. RSD <50%) and then organized by their corresponding transcript profiles into 6 subclusters.

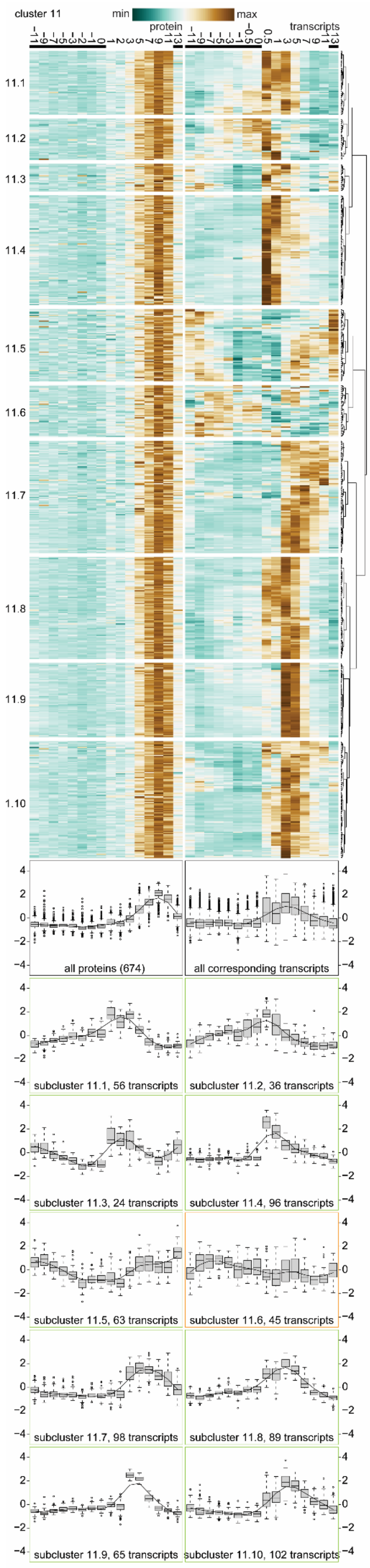


**Supplemental Figure 14:** Cluster 11 of 16, protein abundances peaking ~9 h of the day and associated transcript profiles. Hierarchical clustering was performed first on protein abundances of proteins identified in every time point (av. RSD <50%) and then organized by their corresponding transcript profiles into 10 subclusters.

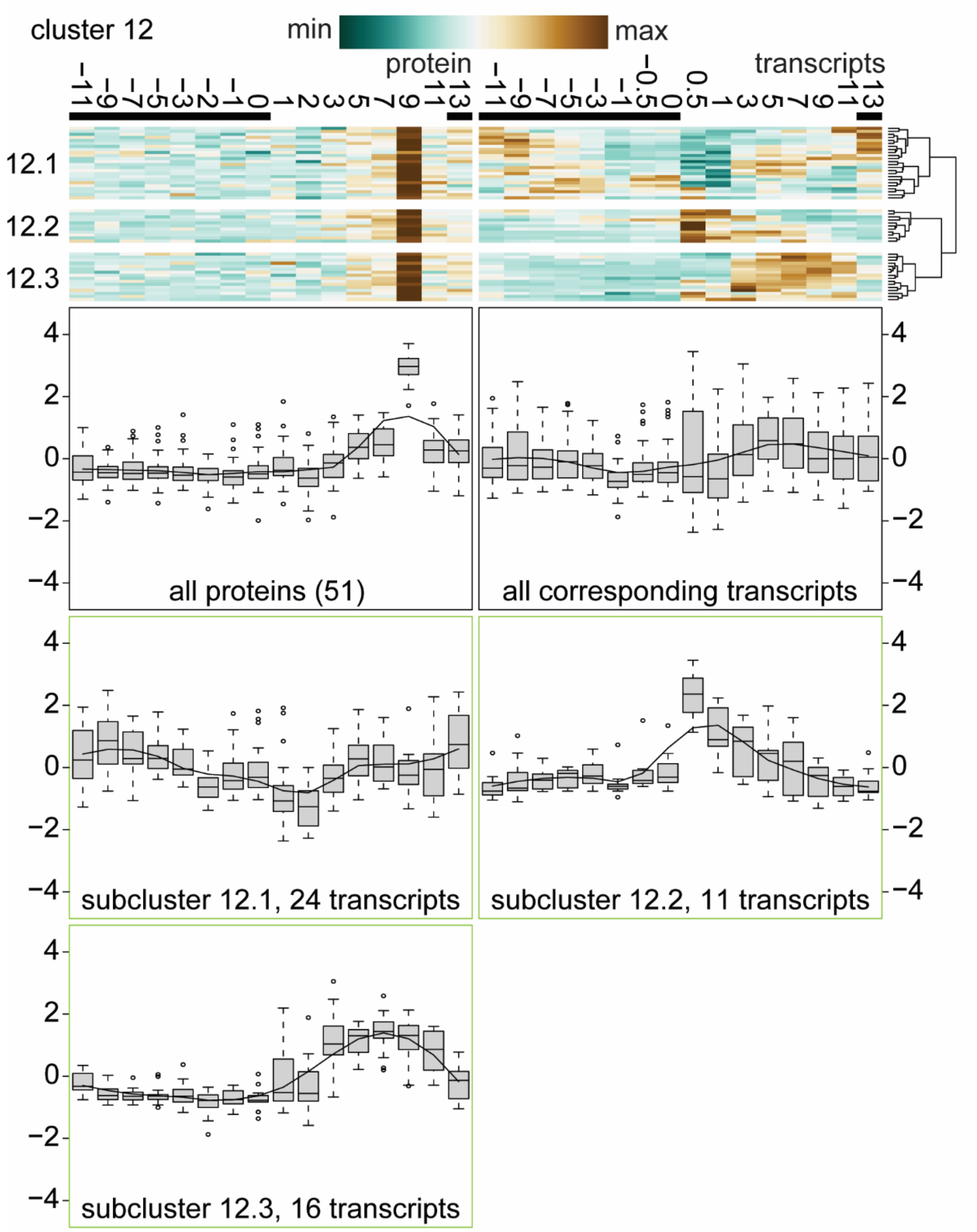


**Supplemental Figure 15:** Cluster 12 of 16, protein abundances transiently peaking ~9 h of the day and associated transcript profiles. Hierarchical clustering was performed first on protein abundances of proteins identified in every time point (av. RSD <50%) and then organized by their corresponding transcript profiles into 3 subclusters.

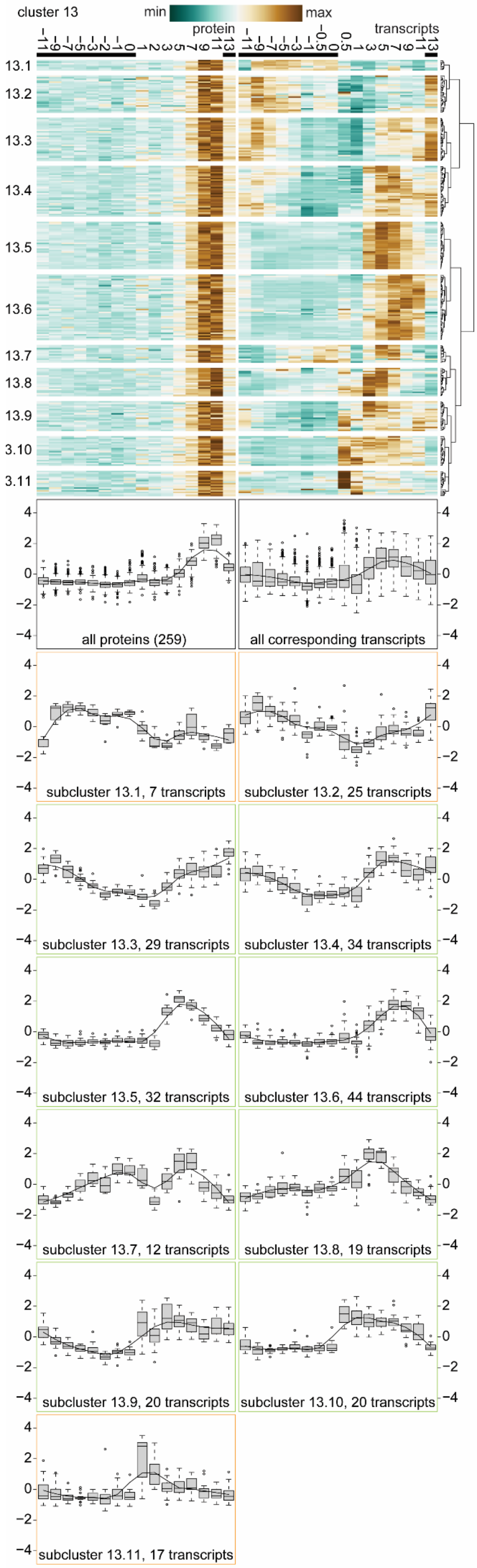


**Supplemental Figure 16:** Cluster 13 of 16, protein abundances transiently peaking ~9-11 h of the day and associated transcript profiles. Hierarchical clustering was performed first on protein abundances of proteins identified in every time point (av. RSD <50%) and then organized by their corresponding transcript profiles into 11 subclusters.

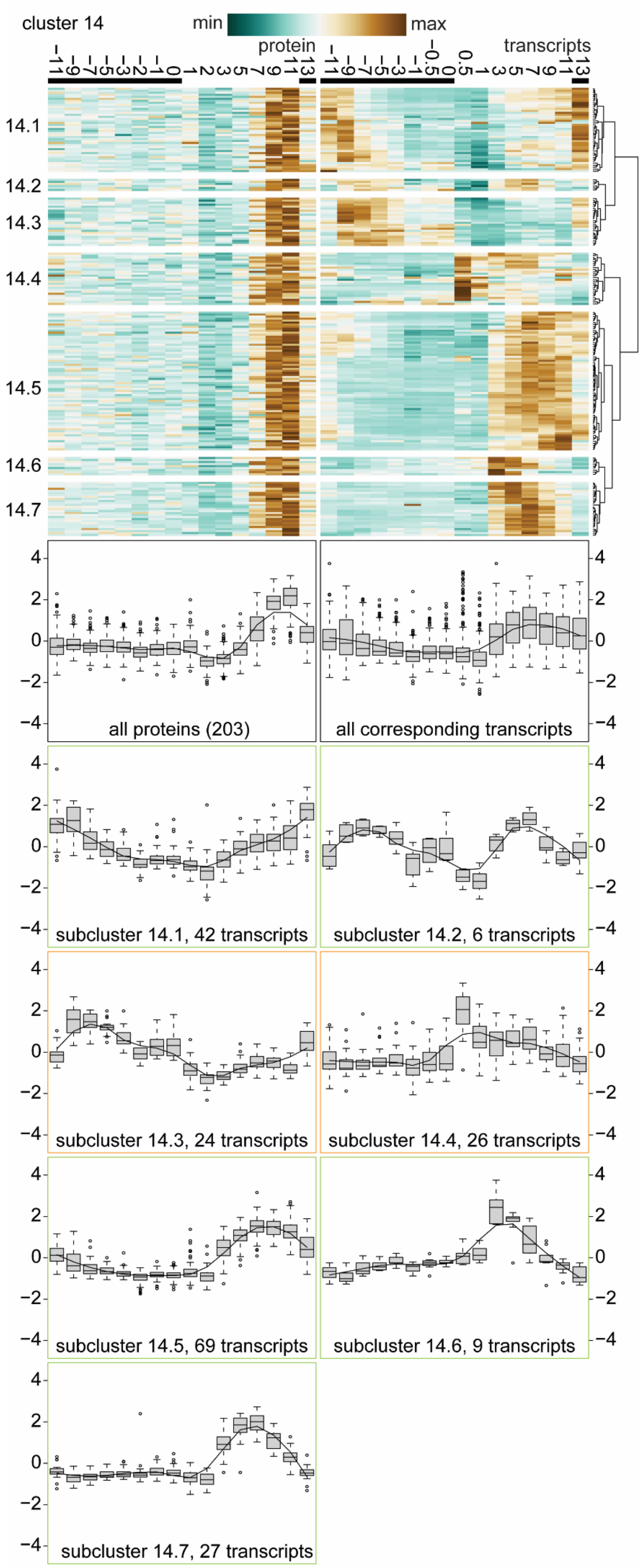


**Supplemental Figure 17:** Cluster 14 of 16, protein abundances transiently peaking ~9-11 h of the day and associated transcript profiles. Hierarchical clustering was performed first on protein abundances of proteins identified in every time point (av. RSD <50%) and then organized by their corresponding transcript profiles into 7 subclusters.

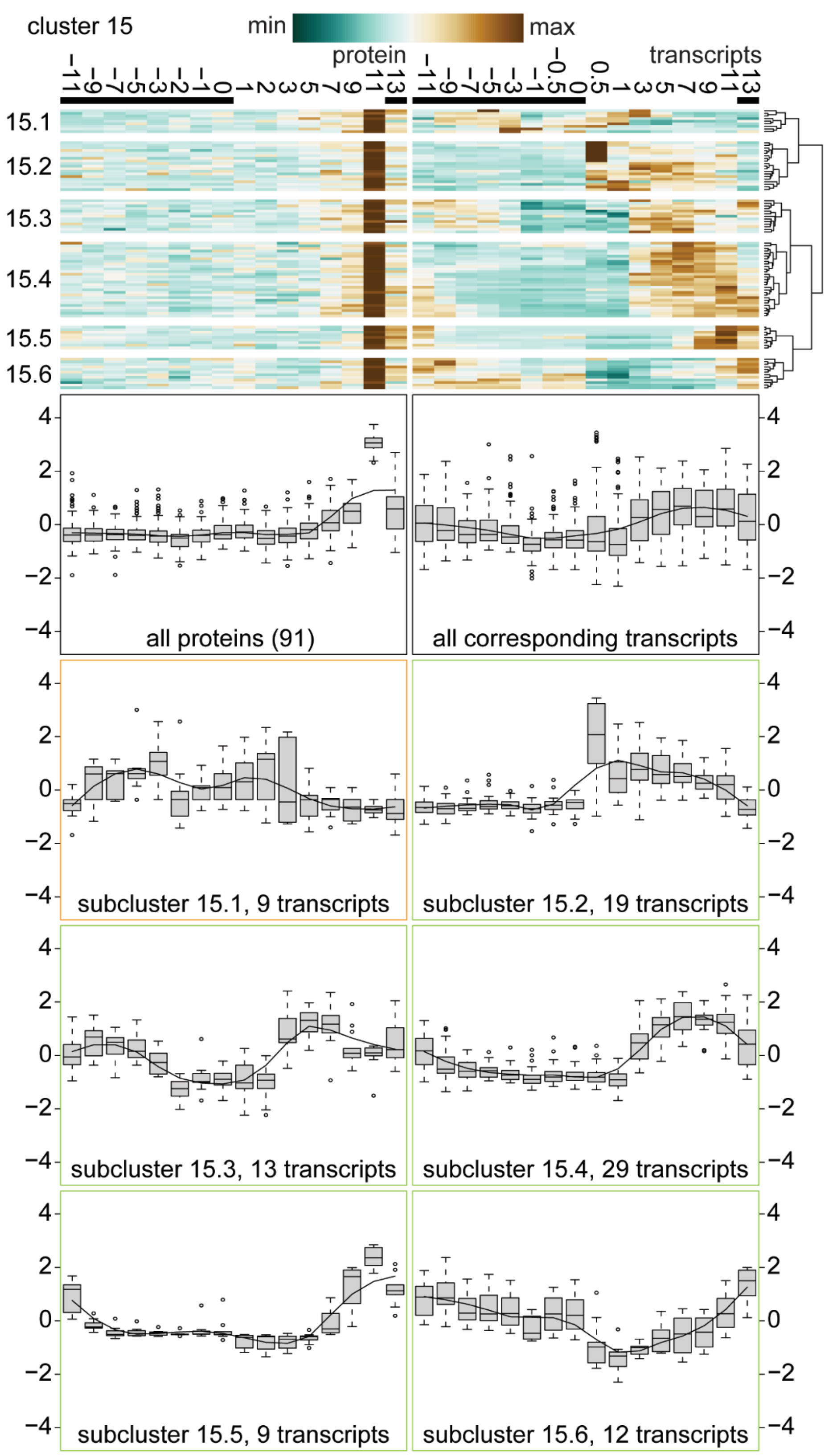


**Supplemental Figure 18:** Cluster 15 of 16, protein abundances transiently peaking ~11 h of the day and associated transcript profiles. Hierarchical clustering was performed first on protein abundances of proteins identified in every time point (av. RSD <50%) and then organized by their corresponding transcript profiles into 6 subclusters.

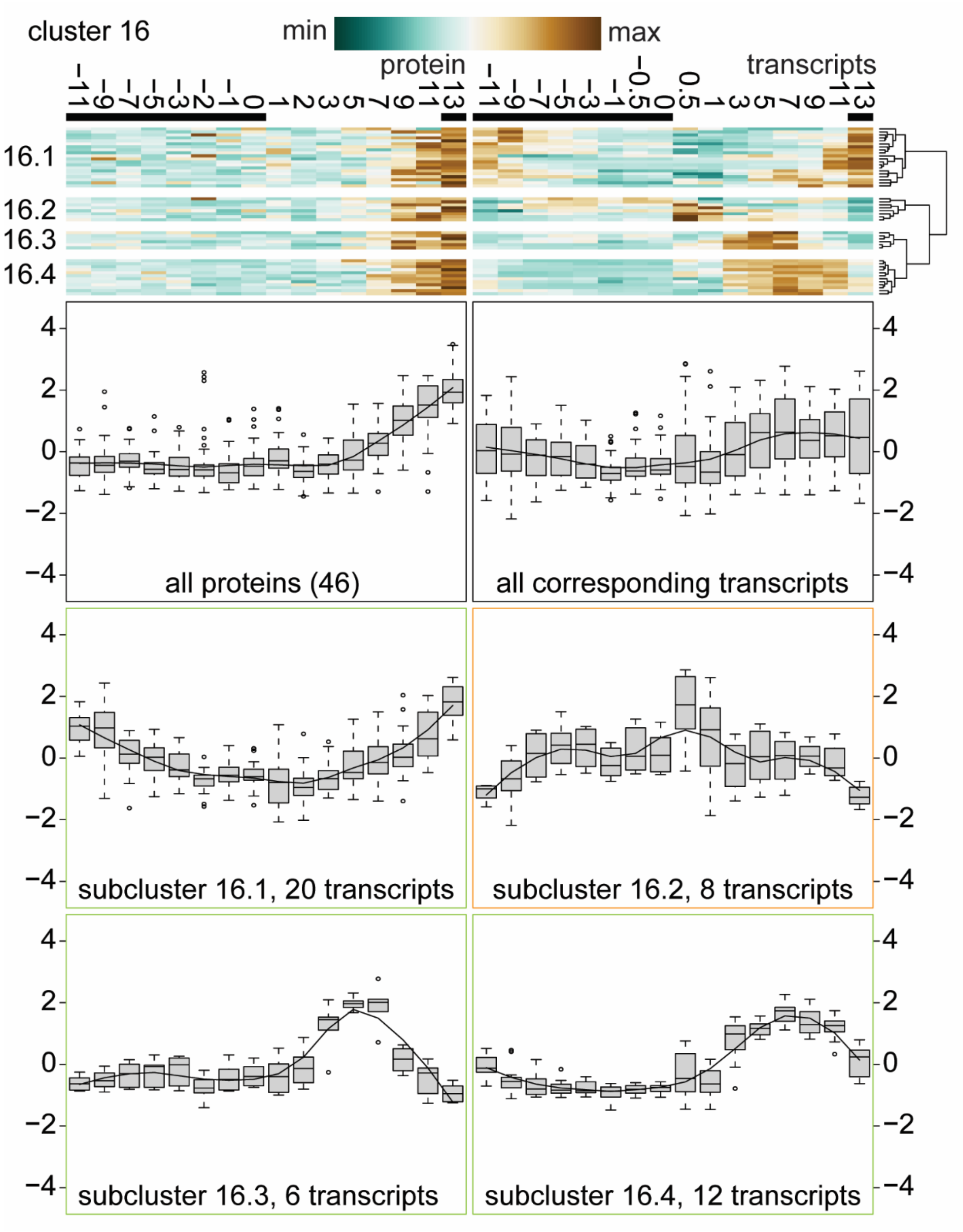


**Supplemental Figure 19:** Cluster 16 of 16, protein abundances transiently peaking late in the day / early night and associated transcript profiles. Hierarchical clustering was performed first on protein abundances of proteins identified in every time point (av. RSD <50%) and then organized by their corresponding transcript profiles into 4 subclusters.
